# Restriction site associated DNA sequencing for tumour mutation burden estimation and mutation signature analysis

**DOI:** 10.1101/2023.07.27.550611

**Authors:** Conor F McGuinness, Michael A Black, Anita K Dunbier

## Abstract

Genome-wide measures of genetic disruption such as tumour mutation burden (TMB) and mutation signatures are emerging as useful biomarkers to stratify patients for treatment. Clinicians commonly use cancer gene panels for tumour mutation burden estimation, and whole genome sequencing is the gold standard for mutation signature analysis. However, the accuracy and cost associated with these assays limits their utility at scale. Using *in silico* library simulations we demonstrate that restriction enzyme associated DNA sequencing (RADseq) may be a cost-effective solution to improve accuracy of TMB estimation and derivation of mutation profiles when compared to a derived FDA approved cancer gene panel TMB score. Using simulated immune checkpoint blockade (ICB) trials, we show that inaccurate tumour mutation burden estimation leads to a reduction in power for deriving an optimal TMB cutoff to stratify patients for immune checkpoint blockade treatment. Additionally, prioritisation of APOBEC hypermutated tumours in these trials optimises TMB cutoff determination for breast cancer. Finally, the utility of RADseq in an experimental setting is also demonstrated, based on characterisation of an APOBEC mutation signature in an *APOBEC3A* transfected mouse cell line. In conclusion, our work demonstrates that RADseq has the potential to be used as a cost-effective, accurate solution for TMB estimation and mutation signature analysis by both clinicians and basic researchers.

## Introduction

Over the past decade, precision medicine has played an increasingly important role in cancer medicine^1^. To identify molecular characteristics which predict response to treatment, high thoughput sequencing is frequently used to comprehensively profile a tumour’s genome. Tumour mutation burden (TMB), a measure of the number of alterations in a cancer genome relative to matched germline from the same patient, has been approved for use as a biomarker to indicate whether patients with any solid tumour type respond to α-PD1 blockade^2^. However, the predictive significance of this biomarker is disputed. While early studies indicated that patients with high TMB were more likely to respond to therapy, many trials have failed to replicate this association^3^.

Extensive analysis of tumour genomic data has revealed that evidence of mutagenic activity in tumour evolution can be derived by categorising mutations into their genomic context^4^. This has led to the development of mutational signatures that reveal how tumours acquire mutations through time. For example, a signature characteristic of UV based mutation can be observed in melanoma samples^5^. This analysis has also shown that breast tumours with a high TMB often have a dominant signature TpCpN>Tp(T/G/A)pN, believed to be caused by activity of APOBEC3A/B enzymes^6–8^. While only a portion of tumours harbour a TMB greater than 10 mutations/megabase, APOBEC hypermutation is overrepresented in these high mutation burden samples^9^. Other common signatures include clock-like processes (COSMIC single base signature (SBS) 1 and 5) and *BRCA* signatures (COSMIC SBS 3 and 8)^10^. Notably, some of these signatures have clinical utility with *BRCA* deficiency identifying patients likely to respond to PARP inhibition. Measuring *BRCA* deficiency through the *BRCA* mutation signature rather than the more conventional identification of genetic mutations in *BRCA1/2* bypasses loss of sensitivity to detect cryptic *BRCA* deficiency (e.g., epigenetic silencing, variants of unknown significance)^11^. Similarly, APOBEC signatures may be a better predictor of response to immune checkpoint blockade (ICB) than TMB alone, or may be a biomarker for Ataxia telangiectasia and Rad3 related (ATR) inhibition^12–14^ thus highlighting the potential of mutation signatures in clinical diagnostics.

Uptake and usage of these “genome-wide” biomarkers in the clinic will be dependent on the availability of inexpensive, simple methods to generate this information. Currently, whole genome sequencing (WGS) is the gold standard approach for elucidating mutation signatures. However, WGS is expensive, requires large amounts of compute and storage power, and many medical facilities do not have access to WGS services. Less complex, targeted genome panels are now routinely used to capture mutation data for relevant tumour suppressor and oncogenes. The FoundationOne Cancer diagnostics panel targets 1.5mbp of exonic tumour DNA, and is the FDA approved companion test for acquisition of TMB as a biomarker for ICB in solid tumour types^2^.

Comparisons of TMB estimates from WGS to those from targeted panels suggest that their accuracy is poor for samples that do not have high TMB^15^ and using other parameters to estimate measurement error via the FoundationOne panel has demonstrated poor results^16^. Additionally, there is substantial variability in TMB estimates derived from the same sample across different labs^17^. The effect of these measurement errors on the capacity to detect associations between response to ICB and TMB has not been formally evaluated. As predictive thresholds have been approved under the assumption that these panels can accurately represent TMB, their ability to correctly identify responders with TMB around the cut-off levels is likely to be limited. Consequently, there is a need for a reliable, accurate, simple and cheap method to estimate TMB which would enable re-evaluation of the prognostic significance of this biomarker.

Restriction site associated DNA sequencing (RADseq) is a reduced representation method used frequently in population genetics for association studies and linkage mapping^18^. RADseq requires a simple library preparation protocol and can be peformed at a low cost per sample. Recently, RADseq based approaches have been used to capture genome-wide mutation data in bladder cancer with RADseq effectively recapitulating mutation signatures in patient derived FFPE blocks^19^. Restriction enzyme based approaches therefore potentially represent powerful tools to supplement targeted sequencing approaches in both clinical and research contexts.

In this paper, we investigated the number of detected mutations required for accurate recapitulation of mutation signatures in breast cancer, and explored the effects of low mutation complexity caused by APOBEC mutagenesis on detection capacity. We then simulated RADseq library preparation methods for breast cancer patients and calculated the accuracy of these methods in acquisition of genome wide biomarkers such as TMB and mutation signatures, benchmarking against our estimate of the FoundationOne Cancer diagnostics panel TMB score. We also simulated use of these libraries to evaluate the effect of TMB on response to ICB. Finally, we demonstrated the utility of RADseq based approaches to study genetic heterogeneity and ongoing mutational processes in a mouse model of breast cancer.

## Materials and methods

### Data download and random mutation subsetting

To randomly subsample different numbers of mutations, WGS data from 560 breast cancer samples, generated by Nik-Zainal et al (2012) was downloaded from Wellcome Sanger Institute ftp website (ftp://ftp.sanger.ac.uk/pub/cancer/Nik-ZainalEtAl-560BreastGenomes/Caveman_560_20Nov14_clean.txt). Data was imported into R (version 4.1.2)^20^, and converted to a gRanges object for compatibility with the SomaticSignatures^21^ R package. Random subsamples of between 10 and 800 mutations were extracted for each sample. Each random subsample catalogue of mutations was then characterised by the 96 trinucleotide context as using SomaticSignatures *mutationContext* and *motifMatrix* functions. This process was repeated times for each mutation subset in each sample to generate 10 representative libraries for each sample at each number of randomly subset mutations.

### Analysis of mutation signature similarities

Mutation profiles were compared by calculating the cosine similarity between the subsetted mutation profile and the WGS profile for each sample using the mutationalPatterns^22^ R package for each iteration and taking the diagonal values of the resulting matrix. The probability of obtaining a cosine similarity to WGS profile of at least 0.9, Pr(CS>0.9), was calculated by dividing the number of cosine similarities for each mutation subset in each sample *s* by 10 (the number of randomly generated libraries for each sample).

In order to formally define the number of mutations required to accurately recapitulate the whole genome mutation signature, a binomial test was used. This was calculated by counting the number of samples that had a >90% cosine similarity to the original WGS profile at each number of mutations subset.

### Defining APOBEC hypermutated samples

APOBEC hypermutated samples were defined as a sample containing >40% Tp(C>T/G)pW mutations in the 96 trinucleotide profile^23^.

### Restriction enzyme mapping

Restriction enzyme information was obtained from the NEB website (https://international.neb.com/tools-and-resources/usage-guidelines/nebuffer-performance-chart-with-restriction-enzymes). Restriction site breakpoints were mapped by scanning the human genome (hg19) for the recognition sites, and defining regions between the breakpoints in a bed-like format. Fragments were restricted to those between 70 and 378 bp commonly used for RADseq experiments. Enzymes that are methylation sensitive were removed due to the fact that different methylation patterns in different samples could result in less reproducible library preparations. Duplicate enzymes (i.e., those with a high fidelity version) were removed. Libraries which returned an average of less than 2 mutations per library/sample were removed. Libraries with greater than 10% coverage of the whole genome were removed as these would not represent practically useful libraries when compared to WGS. The FoundationOne regions were defined by downloading available gene information from the FoundationOne panel^24^ and matching gene names to annotations in hg19. Fragments were restricted to exonic regions as described on the FoundationOne website: for simplicity, we did not include the small amount of intronic regions that are included in the FoundationOne assay. Library coverage was defined using the “reduce” function from the GenomicRanges R package, which gives the expected number of basepairs captured by each library.

### In *silico* library analysis

Once fragments were defined for enzyme libraries and the FoundationOne panel, mutations from the Nik-Zainal dataset were overlapped and cosine similarity was computed for each library as above, using the mutations that overlapped for each library instead of random subsetting. Mutation signature analysis was carried out for each library as decribed above. Mutation rates for each library were defined by dividing the number of mutations overlapping the library by the library coverage, that is, the number of basepairs covered by the library. Mutation rates were then compared with the WGS mutation rate using three methods; Pearson correlation of the WGS mutation library to the mutation rate estimated for the library; and two error-based methods: the difference between the WGS mutation rate and the library mutation rate; and the estimated bias, θ, of the library mutation for the 560 tumour samples. For the error-based methods, libraries with a Pearson correlation value less than 0.9 were removed and a linear model was defined using log2 transformed values of the median value for each library as the dependent variable and log2 transformed values of the library coverage methods as the independent variable in a linear regression analysis.

### Simulated ICB trials

For each sample in the Nik-Zainal dataset, the probability of response to ICB was defined as a function of the WGS mutation rate.The probability was scaled to defined minimum and maximum of 0 and 0.5. This resulted in a value between 0 and 1 for each sample that were that samples probability of response in that trial. The response to therapy in each separate trial was then defined using this probability value. The odds ratio of response vs non-response at defined WGS mutation rate cutoffs was then calculated. For comparison of different library methods used to estimate TMB, the odds ratio of response was calculated as above, except the mutation rate used was for the estimated mutation rate of that sample using the defined libraries. To simulate independent clinical trials where statistics are computed as part of each trial, odds ratio p-values for each cutoff were Bonferroni corrected using *p.adjust* separately for each trial.

This process was repeated 100 times to represent 100 independent trials on the same cohorts of patients. As the rbinom function generates random deviates of the “response” outcome based on tumour mutation burden, each sample was represented 100 times, however the number of trials in which the sample “responded” was dependent on the TMB as encoded in the model. The proportion of trials reporting a significant association at each cutoff was taken as the number of adjusted p-values < 0.05 at each cutoff.

### SSM3 cell culture

SSM3 cells^25^ were cultured in 90% DMEM + 10% FBS, 50μM β-mercaptoethanol, 5μg/mL bovine insulin, 10ng/mL transferrin, 0.3μM hydrocortisone at 8% CO2. Monoclonal isolates were derived by limiting dilution. Following this, a second round of monoclonal isolation was achieved by FACs sorting into 96 well plates. Cells were expanded in 50% conditioned media. Conditioned media was generated by harvesting media from a healthy growing parental SSM3 cell line after 1 day, followed by centrifugation, filtering through a 0.22μm cell strainer and mixing with full SSM3 media in a 1:1 ratio. After initial plating by both methods, single colonies were left to grow for 7 days before replacing conditioned media. Due to the cluster like growth of SSM3 cells, after sufficient expansion of the colony in the 96 well plate, colonies were lifted and replated in the same well in full conditioned media to promote growth and over confluency. After colonies grew to 50% confluency, the entire well was harvested and transferred first to 24 well plates and then to T12.5 and T25 flasks, maintaining cultures in full SSM3 media. Colony growth was heterogenous, however most colonies were able to be transferred after 3 weeks following single cell isolation, and then grown as normal. **Transfection, APOBEC mutagenesis and cell sorting of 0D5 subclones** 0D5 cells were seeded at 800000 cells/well of a 6cm dish. 24 hours after plating, media was replaced with SSM3 media, and cells were transfected with a mixture containing 2μg pcDNA3.1 vectors expressing either APOBEC3A or APOBEC3B (kindly donated by Vincent Caval^26^), co-transfected with 2000μg pcDNA3.1 construct expressing deaminase null APOBEC3A linked to a uracil deglycosylase construct and 1000μg of a plasmid encoding mutant GFP and WT mCherry that is a reporter for APOBEC mutagenesis^27^, with 17.325μL of FuGene in a total volume of 325μL of transfection mixture. Control cells were co-transfected with pcDNA3.1-IRES-GFP^28^ (Addgene plasmid # 51406; http://n2t.net/addgene:51406; RRID:Addgene_51406) and the mCherry reporter construct above. To facilitate setting gates for flow cytometry, separate wells were transfected with 5μg pcDNA3.1-IRES-GFP (GFP+cells), and 5μg of the reporter constructs above (mCherry+ cells), as well as extra wells for unstained and heat killed (Texas Red+) cells. Transfection mixture was left on for 24 hours and then replaced with fresh media. 72 hours after initial transfection, cells were scraped in cold PBS and stained with Texas Red for 20 minutes, washed with PBS and sorted for live, GFP+ cells into 96 well plates containing SSM3 conditioned media on a BD FACS Aria (1:1 ratio of filtered SSM3 media harvested from plates containing 40% confluent SSM3 cells combined with fresh SSM3 media. Single cells were confirmed by phase contrast microscopy, and cells were left to grow for 1 week post sorting. Monoclonal cell lines were then isolated as above (see **SSM3 cell culture**).

### DNA isolation

DNA was isolated from cells by reserving the cell pellet from routine passaging of a T75 cells,. DNA purity was assayed using a Nanodrop spectrophotometer, integrity by gel electrophoresis and quantified by Qubit fluorescence.

### RADseq library preparation and sequencing

RADseq library preparation and sequencing was outsourced to a local company, AgResearch, Dunedin. The RADseq libraries were constructed according to the methods outlined in Elshire et al., (2011)^29^ with modifications as outlined in ^30^. RADseq libraries were prepared using a *PstI-MspI* double-digest, and included negative control samples (no DNA). Libraries underwent a Pippin Prep (SAGE Science, Beverly, Massachusetts, United States) to select fragments in the size range of 220-520 bp (genomic sequence plus 148 bp of adapters). Single-end sequencing (1×101bp) was performed on an Illumina NovaSeq6000 utilizing v1.5 chemistry. Raw fastq files were quality checked using a custom qc pipeline (available at https://github.com/AgResearch/DECONVQC). As one of the qc steps raw fastq files were quality checked using FastQC v0.10.1 (https://www.bioinformatics.babraham.ac.uk/projects/fastqc/).

### Bioinformatic processing of RADseq data and SSM3 WGS data

WGS SSM3 data^31^ were downloaded in FASTQ format from the Sequence Read Archive (SRA) using fasterq-dump from sra-toolkit^32^ (accession SRR2142076). FASTQ files were demultiplexed and adapters trimmed using cutadapt^33^. Poor quality reads from FASTQ files were trimmed using Trimmomatic^34^. Files were then aligned individually using bwa mem with standard parameters^35^. Mutations were called using the GATK4 MuTect2 pipeline for tumour only samples^36^. The MarkDuplicates step was omitted due to the nature of RADseq data likely resulting in a high probability of false positive detection of duplicates, due to reads generated from the same cut site sharing identical mapping coordinates. Sites with <20 reads and an allele frequency (AF) >0.75 were filtered out to remove germline, artifactual and low quality sites. Mutation rate was calculated by counting the number of mutations in each sample and dividing by the total base pairs covered by at least 20 reads (computed using samtools coverage). Cosine similarity matrices were then generated for comparison of different samples. The context of each mutation was extracted using *MutationContext* from the SomaticSignatures package. SigProfiler analysis was carried out using default parameters for extraction of SBS96, with a maximum number of signatures set at 10, and 100 non-negative matrix factorisation (NMF) replicates carried out. The SigProfiler analysis was carried out on samples with each sample split between clonal and subclonal mutations, as well as the SSM3 WGS mutations. Signatures for the computed optimal solution were visually examined and on the basis of similarity, two signatures were combined.

To determine whether samples were enriched for Tp(C>T)pW mutations in the APOBEC transfection experiment, a binomial test was used, with the p-value calculated via

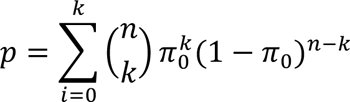

where *k* is the number of mutations in the T(C>T)W mutation context, *n* is the total number of mutations detected in a sample, and *π0* is the predefined proportion of Tp(C>T)pW mutations. For this experiment, *π0* was defined as the proportion of Tp(C>T)pW in the parental 0D5 clone. Thus, if p<0.05 the sample was defined to be to significantly enriched for APOBEC mutagenesis, indicating successful genome editing.

### Code availability

All code and pipelines used in these analyses is available on GitHub (https://github.com/conormcguinness9016/RADSeqPaper).

### Sequence data availability

Demultiplexed RADseq FASTQs and filtered VCFs from monoclonal SSM3 cell lines generated as part of this study have been deposited at the Gene Expression Omnibus (GSE234146).

## Results

### Estimating the number of mutations needed to recapitulate whole genome sequence mutation signatures

To quantify the number of mutations required to recapitulate the whole genome sequence mutation profiles of breast cancers, we used publicly available mutation data from a cohort of 560 breast cancer whole genome cancer catalogues^8^. Random subsets of 10-800 mutations were sampled and a mutation profile representative of the 96 trinucleotide context as described by Alexandrov et al., 2013 was generated for each subsample. This range was used as it represents a range of values that would be captured by reduced representation libraries such as RADseq, up to the number of mutations captured by WGS for breast cancer samples. Each sample was randomly subset as above 10 times. These were then compared to the mutation signatures generated from WGS by calculating cosine similarity. At higher numbers of randomly subset mutations, the cosine similarity to the WGS profile tended to increase (Figure 1a). At n=330 mutations >95% of samples had a cosine similarity >0.9 to the WGS profile (Bonferroni corrected exact binomial test *p*= 4.88×10^5^, 95% confidence interval 95.97-100% of samples with >0.9 cosine similarity, Figure 1b).

**Figure 1.**
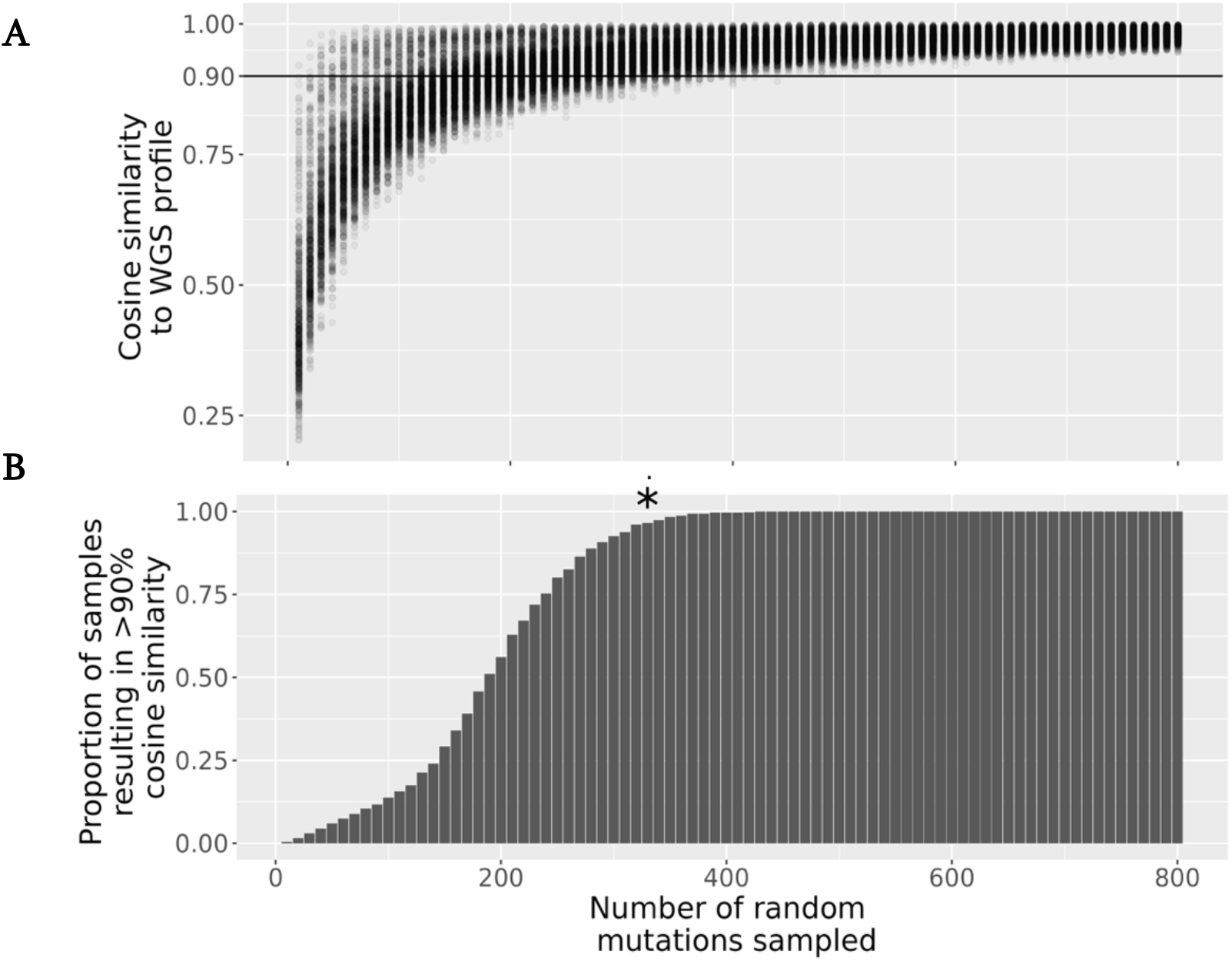
Similarity of mutation profiles derived from WGS and randomly subsetted mutation catalogues from 560 breast cancer profiles.a) Number of mutations sub sampled vs proportion of profiles that result in 90% cosine similarity to WGS profile. Each point is a randomly sampled subset of mutations from one of 560 WGS breast cancer mutation catalogues. b) Number of mutations subsampled vs proportion of profiles that produce a profile with 90% cosine similarity to WGS profile. An asterisk is shown to depict the point at which >95% samples can show >90% cosine similarity to the WGS profile (Bonferroni corrected exact binomial test for proportion of samples >0.95).

### APOBEC hypermutated samples can be reconstructed using a lower number of randomly sampled mutations and represent the majority of high mutation burden breast cancers

A number of samples had a high cosine similarity at a relatively low number of mutations sampled (Figure 1a). We hypothesised that these samples were APOBEC hypermutated samples, as these are characterised by large numbers of C>T mutations in TpCpW contexts, and thus have low mutational complexity. To test this hypothesis, samples were split into “APOBEC hypermutated” or “APOBEC typical” samples (see Methods) and the analysis was repeated as above for the separate groups. APOBEC hypermutated samples had significantly higher cosine similarities compared to APOBEC typical samples (Figure 2a Holm adjusted Wilcoxon rank sum test p<0.05 at each value of mutations subsampled). We repeated the binomial test to define the number of mutations required to recapitulate WGS profiles in these subcategories. APOBEC typical samples were able to be recapitulated at 330 mutations, the same as the global analysis (Figure 2b, Holm adjusted exact binomial test *p*= 0.02, 95% confidence interval 95.58-100% of samples with >0.9 cosine similarity). Conversely, APOBEC hypermutated samples could be recapitulated at 110 mutations (Bonferroni corrected exact binomial test *p*= 3.22×10^-3^, 95% confidence interval 97.33-100% of samples with >0.9 cosine similarity).

**Figure 2.**
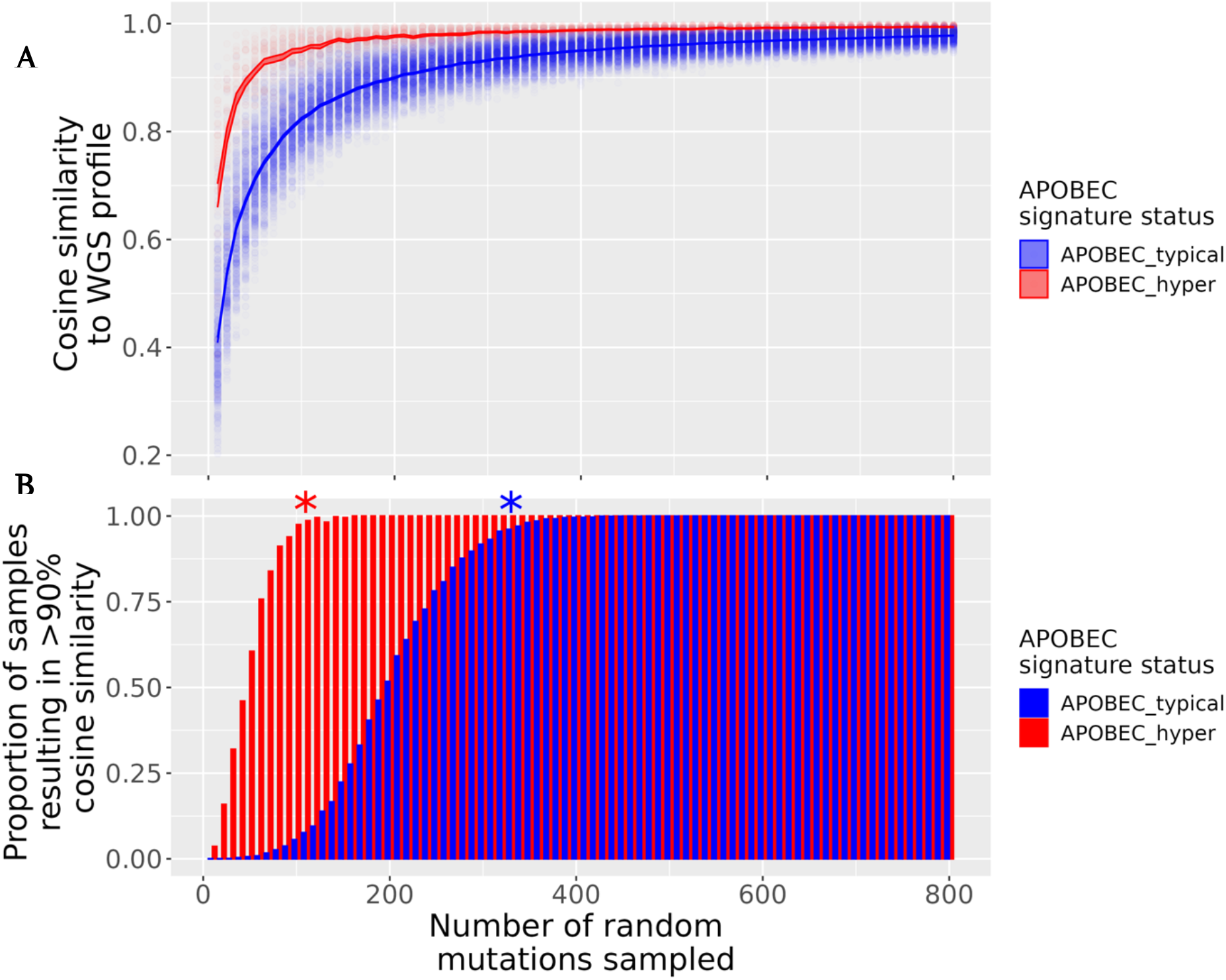
Similarity of mutation profiles derived from WGS and randomly subsetted mutation catalogues from 560 breast cancer profiles. Each sample’s mutation catalogue was randomly subset from 10 to 800 mutations for 10 replicates. a) Distribution of cosine similarity to WGS profiles as a function of number of mutations subset. Each point represents the cosine similarity of a randomly subsetted profile to the WGS profile for that sample. b) number of mutations sub sampled vs proportion of profiles that result in 90% cosine similarity to WGS profile. An asterix is shown to depict the point at which >95% samples can show >90% cosine similarity to the WGS profile (Bonferroni corrected exact binomial test for proportion of samples >0.95) Samples are grouped and coloured by APOBEC hypermutation status, defined as a sample containing >40% Tp(C>T/G)pW mutations in the 96 trinucleotide catalogue.

To examine the distribution of high mutation burden samples and the number of mutations required to recapitulate the WGS profile, samples were colour-coded according to the number of mutations present in the WGS profile. Paradoxically, samples with the highest TMB values were recapitulated at a lower number of mutations, as shown by the darker points on the plot in Figure 3a. Mutation burdens were higher in APOBEC hypermutated samples (median number of mutations=10,110, n=49), than APOBEC typical samples (median number of mutations=3270, n=511) (Figure 3b, Wilcoxon rank-sum *W*=4626, *p*=2.98×10^-13^). When categorised into high (>10 mutations/Mb), medium (5-10mutations/Mb) and low (<5mutations/Mb), APOBEC hypermutation was significantly associated with mutation rate categories (Figure 3b, Fisher’s exact *p*=3.91×10^-11^). When restricted to either “High” (>10 mutations/Mb) or “Not High”, APOBEC hypermutators were significantly more likely to have a tumour mutation burden >10mutations/Mb than APOBEC typical samples (OR=27.77, 95%CI 6.08-172.87, Fisher’s exact p=2.54×10^-6^).

**Figure 3.**
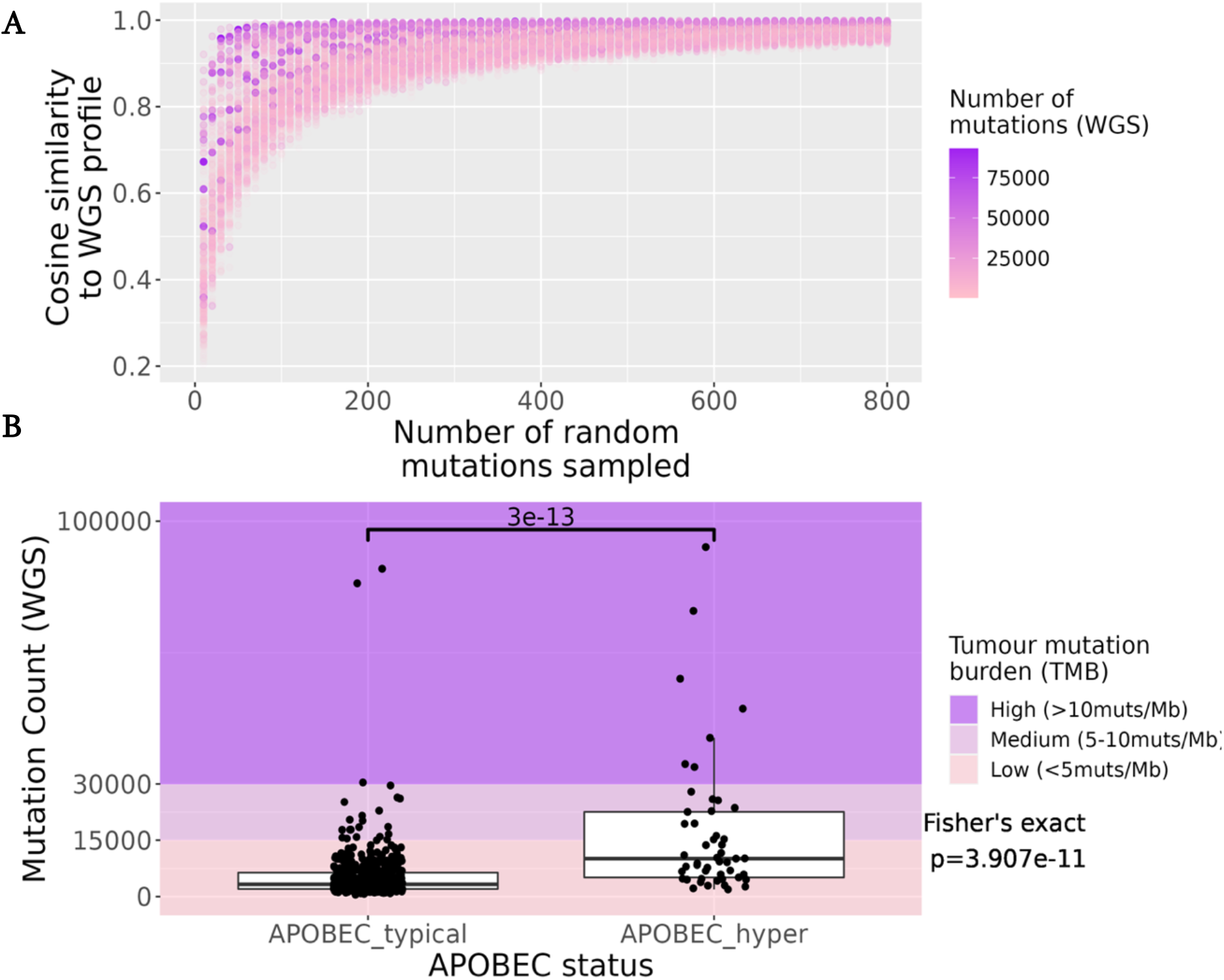
Similarity of mutation profiles derived from whole genome sequencing (WGS) and randomly subsetted mutation catalogues from 560 breast cancer profiles and the distribution of mutation burden between APOBEC hypermutated vs non-hypermutated samples. Each sample’s mutation catalogue was randomly subset from 1 to 800 mutations for 10 replicates. a) Distribution of cosine similarity to WGS profiles as a function of number of mutations subset. Samples are coloured by the number of mutations in the sample. Each point represents the cosine similarity of a randomly subsetted profile to the WGS profile for that sample. b) The distribution of mutation counts between non-APOBEC hypermutated (APOBEC_typical) and APOBEC hypermutated (APOBEC_hyper) samples. The p-value of the Wilcoxon rank sum test between the two groups is shown, as well as the Fisher’s exact p-value to test for association n(bottom right). APOBEC hypermutation status was defined as a sample containing >40% Tp(C>T/G)pW mutations in the 96 trinucleotide catalogue.

### Optimisation of reduced representation methods to capture mutation signatures

The potential of RADseq to recapitulate genome wide mutation characteristics was examined. Furthermore, our estimate of the FoundationOne (F1) cancer diagnostics panel TMB was compared to these methods as it is the FDA approved standard for measuring TMB. We used the same mutation data as described above. However, instead of randomly subsetting mutations, we subset mutations based on whether they would be captured by an in silico genome digest, for each of 277 enzymes. First, we removed “high fidelity” enzymes, as these are just variants of restriction enzymes that recognise the same cut sites, and therefore would result in duplicate libraries. Libraries were removed if the enzyme used was methylation sensitive, or if the median number of mutations captured was less than two. Methylation sensitive enzymes would be less amenable to RADseq on cancer samples as the captured regions would be dependent on methylation at each locus. The genome coverage parameters allowed us to focus on libraries with a high enough coverage to capture mutations for signature analysis.

After filtering, 117 enzyme libraries remained for analysis. For the F1 panel, we restricted mutations to those in exonic regions of genes reported to be assayed by the panel. The libraries captured a range of mutations in the range of our random sampling trials (range 1-54,968 mutations captured, Figure 3.5a, Supplementary Figure 1a). Consistent with our random subsampling results, libraries with lower coverage, such as the F1 panel, and rarely cutting enzymes such as *Kpn*I, usually resulted in lower mean cosine similarities of mutation profiles to the WGS profile while higher coverage restriction enzyme based methods were able to robustly recapitulate all mutation signatures (Figure 4a,b, Supplementary Figure 1b). Interestingly, some libraries did not follow this general trend. For example, while more mutations were captured by the *MspI* library than the lower coverage *BpuEI* library, the cosine similarities to the WGS profiles of these libraries were similar. When samples were split by APOBEC hypermutation rate as above, even low frequency cutters such as *SpeI* were able to accurately recapitulate some APOBEC hypermutated samples (Figure 4d, Supplementary Figure 2). The mutation profiles generated from simulated F1 libraries were highly dissimilar to the WGS generated profiles, with only a portion of profiles in the APOBEC hypermutated cohort showing similarity to the WGS profile (Figure 4c,d).

**Figure 4.**
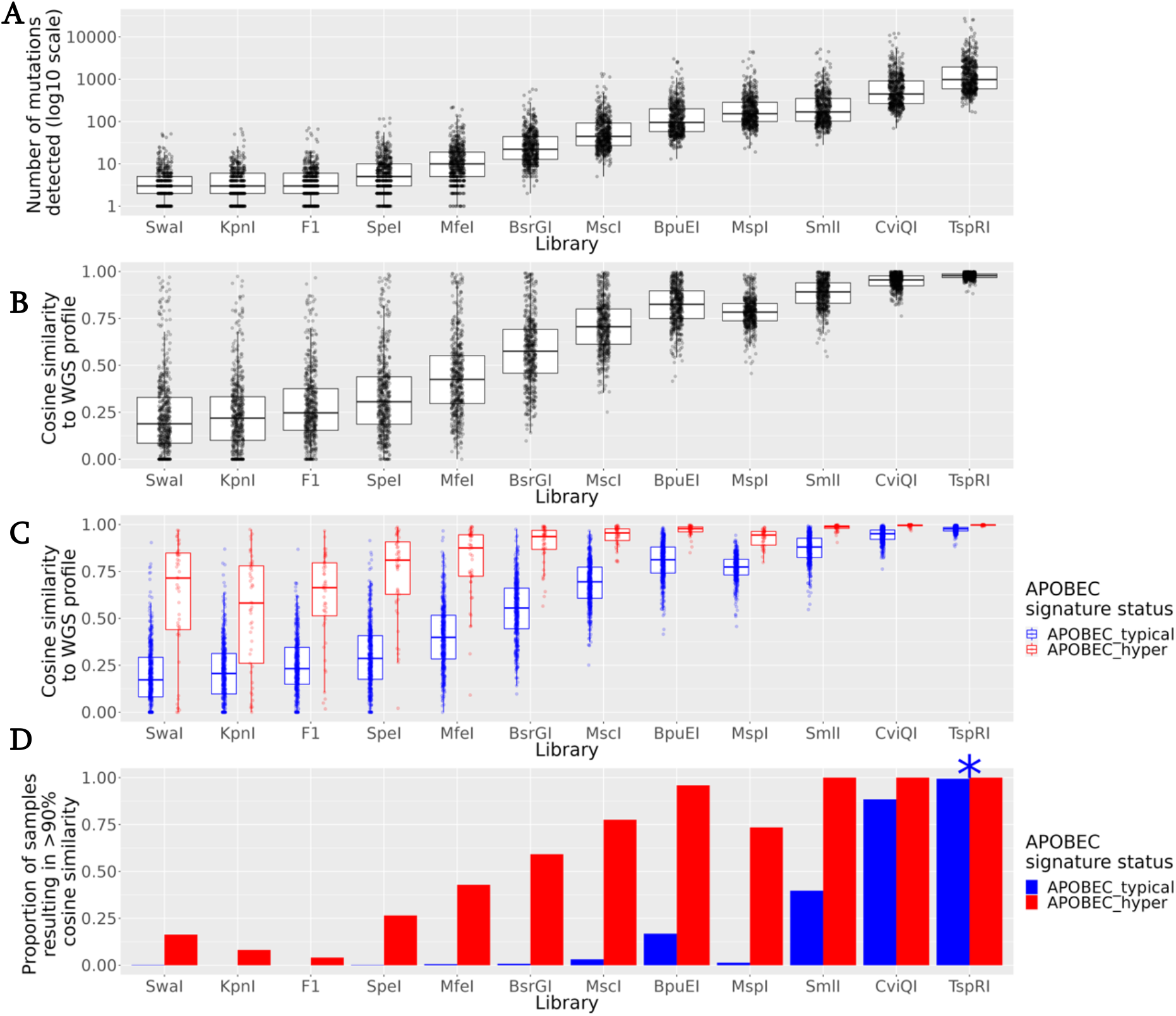
Representative library capture information. a) number of mutations captured by each library for each of 560 breast cancer genomes. b) Distribution of cosine similarity of the library 96 trinucleotide signature profile to the WGS trinucleotide signature profile. c) same as b) but split by APOBEC hypermutation status. d) Library vs proportion of profiles that result in 90% cosine similarity to WGS profile. Samples are coloured by APOBEC hypermutation status as in c).

### Optimisation of reduced representation methods to estimate TMB

The suitability of various library preparation methods to estimate TMB was then tested. Estimation of tumour mutation rate derived from all simulated library preparation methods were significantly correlated with values for the whole genome sequence values (Pearson’s r>0.65, adjusted p<0.05 for all libraries, Figure 3.6a). Evaluation using correlation however, has been criticised, due to the fact that highly mutated values can skew this parameter ^37^. This effect is demonstrated in Figure 5b. While the overall correlation appears to be strong for the F1 library, the wide range of points for values below 10 mutations/Mb in the F1 library suggests that this method cannot reliably estimate TMB. For example, many samples with WGS TMB between 0 and 2.5 mutations/Mb have highly variable estimates of F1 TMB, such that many samples in this category would have an estimated TMB>2.5 mutations/Mb using the F1 library. By comparison, the *SpeI* library, with roughly double the coverage of F1, is both highly correlated, and displays much lower variability, even in samples with lower TMB.

**Figure 5.**
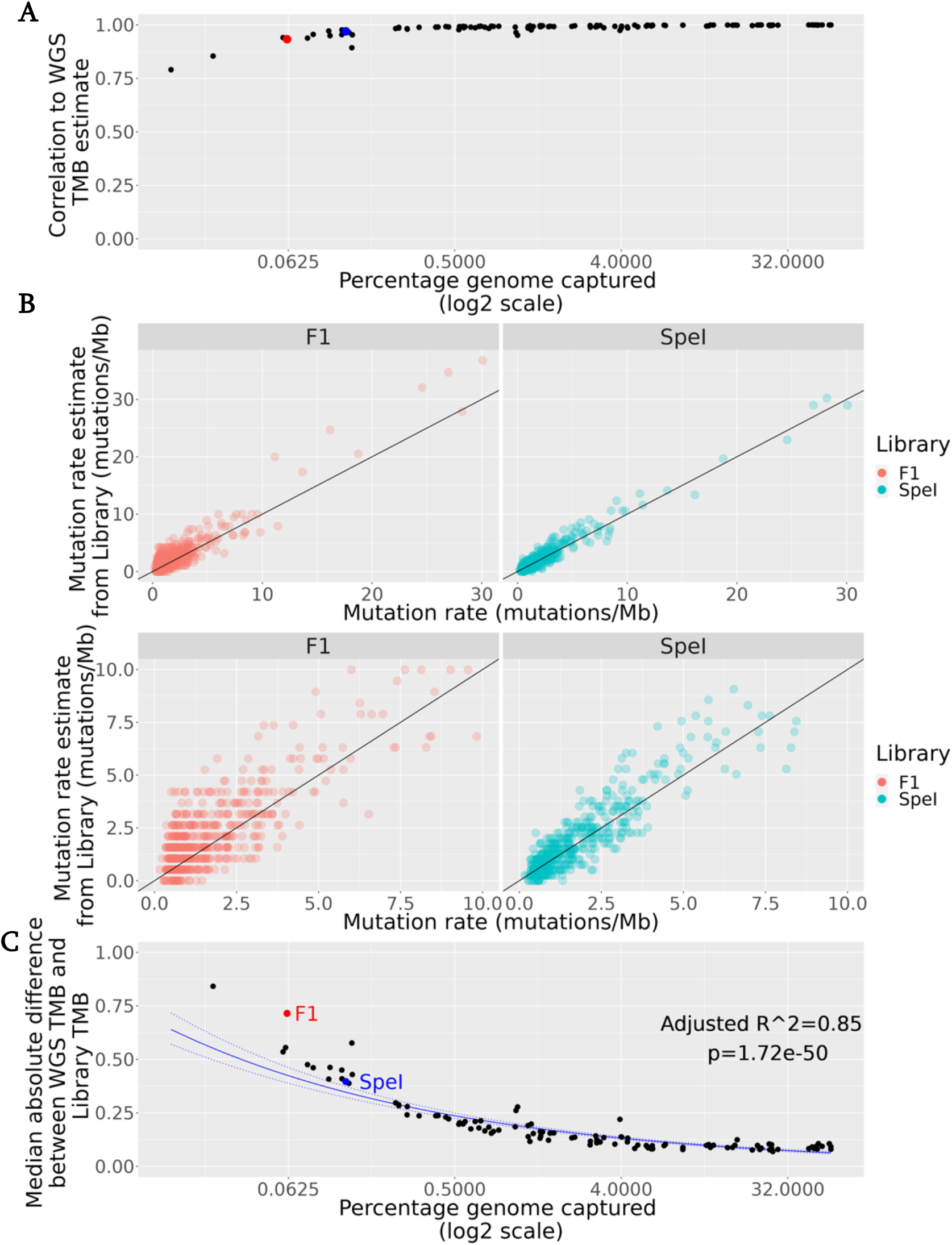
Correlation and error rates of libraries in estimating TMB. a) x axis depicts the coverage of libraries, y axis depicts Pearson correlation values between WGS TMB estimates and library TMB estimates. The F1 and SpeI libraries are highlighted as red and blue points respectively. b) Scatterplot of mutation rate detected in the WGS profile vs the mutation rate detected by the F1 library (left panels, red) or the SpeI library (right panels, blue). The top panels show all the samples, while bottom panels depict only those with a WGS TMB <10 mutations/Mb. c) The median absolute difference of the WGS TMB estimate and the library TMB estimate) (y) as a function of library coverage (x). A regression line (solid blue) is depicted with 95% confidence intervals (dashed blue). The F1 and SpeI libraries are highlighted and labelled in red and blue respectively.

Because of the potential issues with correlation-based assessment, we utilised two other parameters to measure the validity of each library in estimating TMB; the median absolute difference between the library and WGS mutation rate; and the median estimated bias, θ^16^. As expected, as the proportion of the genome covered increased, the median error rate of the library, as estimated by both parameters, decreased exponentially (Figure 5c, Supplementary Figure 3, Supplementary Figure 4). A linear model was fit, and this model suggests that error rates in this regard can be halved by increasing genome coverage by 8-fold. Libraries with a higher rate of error in this regard would be expected to have a median absolute difference higher than the 95% confidence intervals predicted by this model. The F1 library clearly falls outside of the 95% confidence interval of this linear model (Figure 5c). Some libraries with similar coverage, such as the *SpeI* library performed better than the F1 library in this regard (Figure 5c).

### Implications for immunotherapy trials

To examine the potential implications of the above findings in evaluating the significance of TMB as a biomarker for potential response to ICB, we simulated trials of ICB in breast cancer. The model assumes that the probability of response increases moderately as a function of mutation rate, plateauing at a probability of 50% at 11 mutations/Mb (Figure 6a). Although a simple model, this allowed us to examine the probability that, using a defined library preparation method in a cohort of breast cancer patients, the trial could detect an effect at a certain TMB cutoff. First, we tested whether, based on the results of Figure 1, pre-screening for APOBEC hypermutation might increase the power of such trials. In 100 simulated trials, clinical benefit, in the form of a significant odds ratio at a defined TMB cutoff, could be detected in the APOBEC hypermutated cohorts, with just under half the trials achieving a significant association with response (Figure 6b and c). By contrast, unselected cohorts almost always failed to detect any association of mutation rate with response (Figure 6b and c).

**Figure 6.**
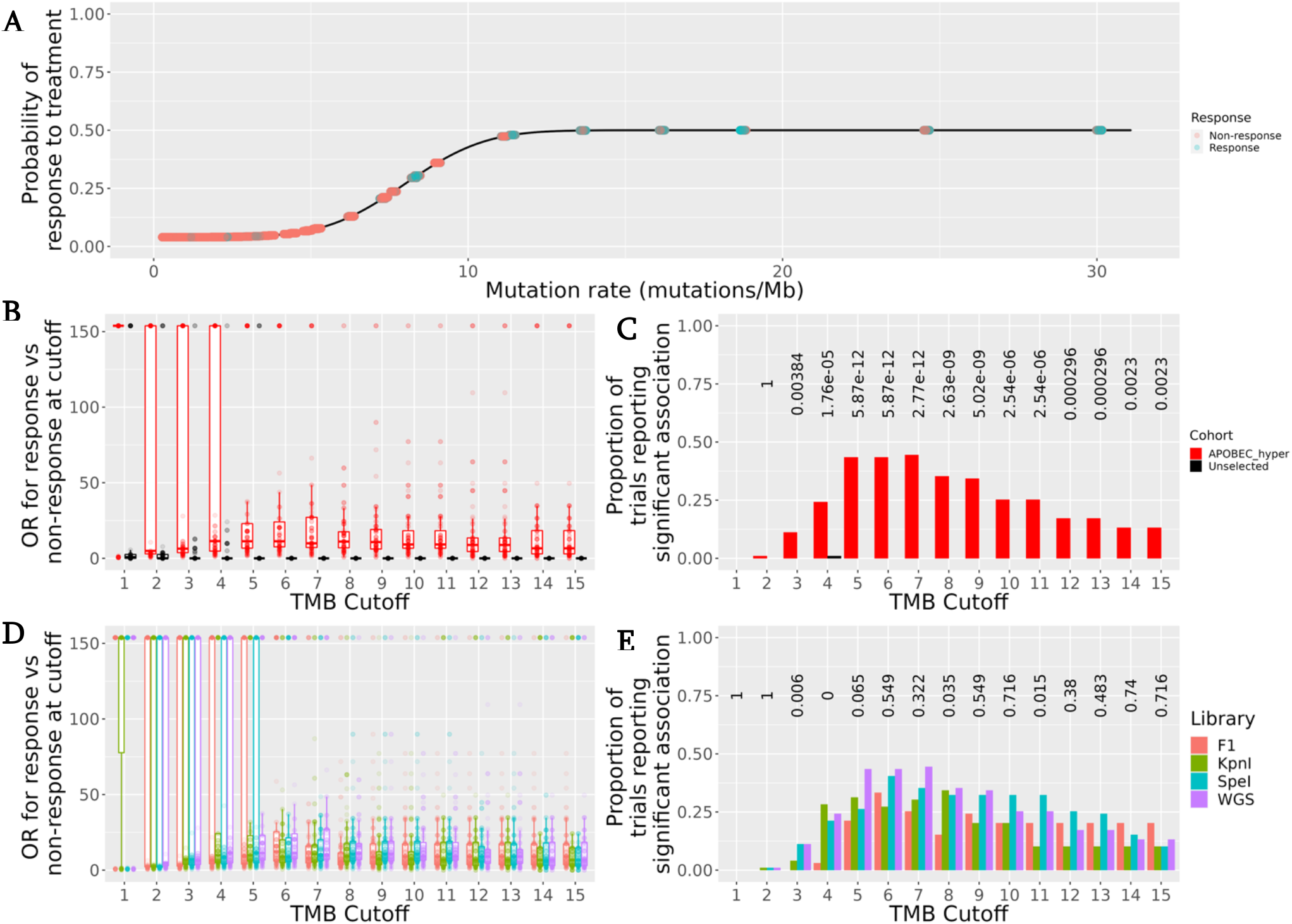
Simulated immune checkpoint blockade trials. a) Model tested, whereby the probability of response to ICB increases as a function of the TMB (mutations/Mb) (x axis). The distribution of the model is depicted in black. Samples tested are depicted on as jittered points, where the response to ICB is depicted as the colour. b,d) odds ratio for response (y) at defined for defined mutation cutoff (x) in each of 100 different trials (depicted as points). Trials are coloured either by cohort b) or by library used to estimate TMB. c,e) Proportion of trials reporting significant association (y) at a defined cutoff (x). Bars are coloured as in b) and d). Above are Fisher’s exact test p-values testing proportions reporting significant association bv cohort (c) or by library used to measure mutation rate (d).

Next, we tested whether the library used to detect this association could affect the results of these trials. To do so, we computed significance as above, using the APOBEC hypermutated cohort, however we used the mutation rate estimated by the library to test the association, instead of the ground truth values derived by WGS. The proportion of trials reporting a true association tended to be lower when using the F1 panel at all cutoffs (Figure 6d,e) compared to using a WGS value. This trend remained true when using the widely used cutoff of 10 mutations/Mb. To test whether RADseq libraries could outperform F1 in this respect, additional libraries were tested. We used the *KpnI* library, as it has a similar library coverage to the F1 library and the *SpeI* library, which has roughly 2x the coverage of F1. At low TMB cutoff values (e.g. between two and four mutations/Mb), where use of F1 would result in no association being reported, use of *KpnI* libraries was able to detect a statistical association. The *SpeI* library was consistently able to improve the loss of significance seen at higher cutoffs (from 6-10 mutations/Mb).

### Utility of RADseq based methods to track mutation signature changes in a mouse cell line

While many mutation signatures have been defined using existing sequencing data, the processes by which these signatures arise remains largely unknown. To empirically test the ability of RADseq to detect mutation signatures and estimate TMB, we derived monoclones of the SSM3 cell line and subjected them to RADseq, then compared the resulting profiles to published WGS data^31^.

To determine whether a known mutation signature can be recapitulated from RADseq data, we transfected the 0D5 cell line with plasmids encoding either *APOBEC3A* and *APOBEC3B,* along with a uracil deglycosylase inhibitor (UGI) construct and a reporter construct that encodes a mutant GFP that is reactivated upon APOBEC editing^27^. As a control, we cotransfected a population with plasmids encoding WT GFP and mCherry and sorted double positive cells. Derivative cell lines were then sequenced using a RADseq protocol and compared to published WGS sequencing data. When compared to the published WGS data for the SSM3 cells, all clones, including the parental 0D5 clone, had increased mutation rate when measured by RADseq (Figure 7a). Several clones derived from the population transfected with the *APOBEC3A* constructs had an increased mutation rate, and one clone, A2H1, had a different mutation profile to the remainder of the clones characterised (Figure 7a,b). Inspection of the 96 trinucleotide mutation frequency of this clone revealed that A2H1 had a significantly increased rate of mutations in a T(C>T)W context (Figure 7c), the preferred substrate for APOBEC enzymes. Furthermore, the mutations in the T(C>T)W contexts were clearly enriched in the YTCA context, supporting the notion that APOBEC3A mutagenesis generated these additional mutations (Supplementary Figure 5).

**Figure 7.**
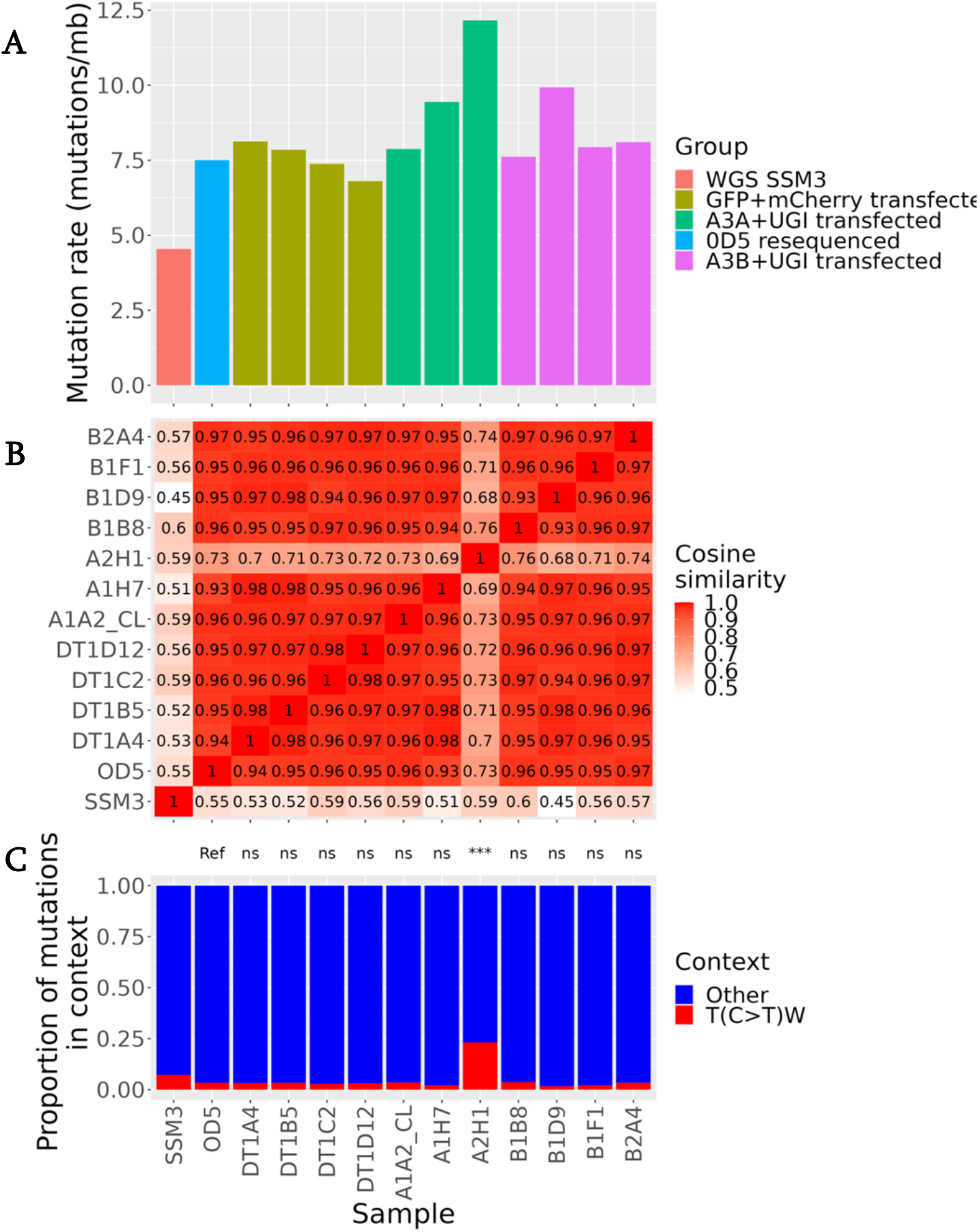
Genomic profiling of derivative 0D5 clones transfected with APOBEC mRNAs using RADseq. a) mutation rate (mutations/Mb sequence) in different samples. b) Cosine similarity of mutation profiles. Tiles are coloured by value, with the cosine similarity value between two profiles printed inside each tile. c) Proportion of mutations in either T(C>T)W or other contexts in each sample. Above, results of one-sided proportion test for proportion of T(C>T)W mutations in each sample, using the proportion in the 0D5 parental clone as a reference. ns=not significant, ***=p<0.001

## Discussion

Genome wide parameters such as TMB and mutational profiles have the potential to play a significant role in precision medicine, either as biomarkers, or as an aid in understanding the aetiology of cancer mutagenesis. We used our own data, together with previously published data, to explore alternative approaches for the assessment of these parameters and investigated the potential implications of the methodology used for cancer immunotherapy research. We identified that, based on random subsampling of mutation catalogues, mutation profiles could be accurately reconstructed for most tumours in the cohort using 330 mutations. APOBEC hypermutated profiles, presumably due to their low mutational complexity, were able to be recapitulated at 110 mutations. This finding may have important clinical ramifications, as it implies that these tumours can be effectively characterised with high confidence from lower coverage panels or via methods such as sequencing of ctDNA<u>^1,2^</u>. Additionally, in this small cohort, APOBEC hypermutators were significantly more likely to have a mutation burden greater than 10 mutations/Mb when compared to non-hypermutators, consistent with previous reports^9,39^. The utility of RADseq to reconstruct mutational profiles and estimate TMB was evaluated and compared to an estimate of the TMB from using the genes analysed in the FDA approved F1 panel. While TMB estimation in this cohort was significantly correlated, libraries of similar coverage outperformed the F1 panel in terms of error and potential utility in simulated ICB trials. We suggest that the current use of the F1 panel, combined with the use of unselected cohorts (compared to potential APOBEC hypermutators) significantly reduces the power to determine whether TMB could be a predictive biomarker for response to ICB. Finally, we demonstrated the utility of RADseq to detect changes in mutation profiles in mouse cell lines, including detection of an APOBEC hypermutation signature.

According to our simulated ICB trials, current clinical trials are likely to be underpowered to truly answer whether TMB can be used as a predictive biomarker for ICB. This may be because unselected cohorts do not represent the full spectrum of TMB in breast cancer. Therefore, pre-selecting APOBEC hypermutated samples may enable assessment of whether there is a significant association between TMB and ICB response. Previously, hypermutation of COSMIC signatures 2 and 13 in breast cancer has been associated with the deletion polymorphism affecting the *APOBEC3B* locus, especially in ER+ breast cancer (Nik-Zainal *et al.*, 2014; Pan *et al.*, 2021). The frequency of this polymorphism varies based on ancestry. For example, while the MAF is low in European and African populations, East Asian populations exhibit heightened frequency ^41^. In carriers of the polymorphism, a high TMB is associated with increased immune infiltration and enrichment of an adaptive immune response signature^13, 14, 39, 40^. In addition, in other cancer types such as lung cancer and cervical cancer, APOBEC hypermutation has been associated with these characteristics and improved response to immune checkpoint inhibition, even in samples that did not pass the “high TMB” threshold of 10 mutations/Mb^12–14, 42–45^. This suggests that it is possible that APOBEC hypermutation, may in itself be predictive of response to immunotherapy. Therefore, an APOBEC enrichment score, combined with a TMB estimate, may provide additional clinical utility compared to a TMB estimate alone.

Our data also suggest that use of the FoundationOne panel to evaluate TMB as a predictive biomarker in immunotherapy trials may reduce the power of these trials. Detection of hypermutated samples was accurate and concordant with a similar experiment carried out by FoundationOne, using lung and colorectal TCGA data^46^. An ongoing query in the field, however, is the determination of an optimal cutoff point for high TMB samples. The currently approved threshold for this biomarker is >10 mutations/Mb^2^, which would be an acceptable cutoff using this panel due to the reduction of error above this cutoff. Post-hoc analyses from a range of different tumour types, however, have suggested that optimal cutoffs are likely to differ between tumour types^3,47^. These analyses have suggested that tumour types with lower median TMB such as breast cancer do not demonstrate an association between TMB and response to ICB, in comparison to tumour types with high median TMB. Our results suggest that the use of the F1 panel, and perhaps cancer gene panels in general, have poor accuracy when estimating TMB in these “lower TMB” cancer types. This may be due to the fact that these panels target known cancer genes and therefore, due to selection pressure, alterations would be expected to be found at different frequencies in these genes when compared to the rest of the genome. Therefore, we suggest that the lack of associations seen in these trials could be a result of the measurement error associated with the use of these panels to estimate TMB. Our simulated ICB trials, for example, demonstrated that at lower cutoffs, the inaccuracy of the F1 panels leads to a loss of power to determine whether TMB is predictive of response. As the currently accepted value has been based on a small range of tissue types, exploratory trials in other solid tumour types measuring TMB using this panel should be cautious about concluding associations based on the TMB estimation, especially in tumours with with lower median TMB. If a tissue’s true predictive value is lower than this threshold of accuracy, any true benefit may be lost due to poor estimation below this threshold. Our results suggest that optimising RADseq based approaches to estimate TMB should resolve these issues, as they have less error in TMB estimation than the F1 panel, even at lower coverage.

Another potential source of discrepancy between these results is that the FoundationOne cancer diagnostics panel sequences at a very high depth (>500x), as the panel may detect mutations at very low variant allele frequency (VAF)^1^, and therefore overestimate clonal TMB. As the data used in this study was not of a sufficiently high depth, we were unable to model the extent and consequences of this overestimation of TMB. Tumour diversity may be associated with response to immune checkpoint blockade; genetically heterogeneous tumours tend to elicit a relatively poor response to ICB, in comparison to clonal tumours^49^.Therefore, a high TMB detected by “deep and narrow” sequencing methods may represent a clonally diverse tumour with low numbers of mutations in each clone rather than a single clonal population with a high number of mutations and potential antigens. It is currently unknown how VAF values should be integrated into the TMB report^46^. Paradoxically, “shallow and wide” may be a more appropriate approach as variants that are detected will be present at a high VAF in the tumour. Altogether, these data indicate that for studies into previously uncharacterised tumours undergoing immune checkpoint blockade, estimations of TMB based on panels targeting a small fraction of the genome should be used with caution to determine threshold values that guide clinical decisions of this biomarker.

Comparatively, the simulation of restriction enzyme based methods showed that these methods can achieve high coverage, provide accurate estimates of TMB, and effectively recapitulate genome wide mutation signatures. Although we only considered single enzymes in our analysis, combinations of enzymes may improve coverage; the potential of using combinations of enzymes to recapitulate mutation signatures in oesophageal cancer has been demonstrated^19^. These methods have additional benefits over panel-based approaches, in additioan to increased coverage. For example, library preparation methods are relatively simple, cost-effective, and many labs may already have access to the resources needed, and there are less PCR enrichment steps which may lead to bias in panel-based analysis.

We have demonstrated the potential use of restriction enzyme-associated library preparation methods to ascertain mutation signatures in the murine SSM3 cell line. This is an ER+ breast cancer cell line that is able to be implanted into immunocompetent, syngeneic 129SvEv mice, a potentially powerful model for studying immune populations in ER+ breast cancer, and the effects of endocrine therapies on ER+ tumour biology. Our transfection/cell sorting experiment results demonstrate the utility of RADseq to ascertain the aetiology of mutational processes acting in tumours, specifically the APOBEC mutation signature. Transfection of *APOBEC3A* mRNA resulted in a characteristic APOBEC signature. Our data suggested that the corresponding murine *Apobec3* did not actively generate the APOBEC signature which is supported by the observation that the genetic sequence of this gene shares little homology to mutagenic human APOBEC enzymes^50^, and none of the control/APOBEC3B clones in this experiment demonstrated an increased APOBEC mutation signature. However, we cannot rule out the possibility that RADseq did not detect regions that were exposed to *kataegis* mutational processes, which have been assigned to both APOBEC3A and APOBEC3B activity^51^. Despite this, this experiment demonstrates the high mutagenic potential of this enzyme, even over period of time estimated to be equivalent in this case to a single cell division. Intriguingly, another clone in this group also demonstrated increased mutation rate, albeit without the APOBEC signature. This may have resulted from the of loss of cells that were unable to survive mutagenic stress, leaving only clones with high “mutagenic fitness”, and potentially already high mutation rates, to survive transfection with this mutagenic enzyme.

One of the shortcomings of our study is the lack of prospective testing on new sequence data from patients and empirical comparison of reduced representation methods. We instead relied heavily on a well characterised mutation dataset, assuming that mutations would be captured just as accurately using the reduced representation methods. Previous work has shown concordance between restriction enzyme based methods and whole genome sequencing in ascertainment of mutational profiles and as noted above we reaffirmed this finding in a non-human model^1^. Another potential issue is that restriction enzyme sites themselves can be mutated, potentially obfuscating our *in silico* digest simulations. This may be exploited by researchers, especially if detection of a shift in mutations to an *a priori* known signature is required; for example, researchers may exploit the bias for APOBEC mutagenesis in TpCpA contexts by digesting the genome with *BclI* (recognition sequence TCATGA).

Despite this, we were able to replicate the previously reported utility of restriction enzyme based library preparation methods in both a different tumour type, with different mutation characteristics, and in a non-human model of cancer. Future applications may range from detection of mutation shifts in cell lines akin to experiments outlined by Zou et al. (2018)^53^. These experiments, previously restricted by cost and access for many research labs, can be performed in small labs for a fraction of the price of WGS. Combined with data from Perner *et al.,* (2020), this technique represents a cost effective tool for characterisation of mutation signatures in cancer. Experimental approaches, and larger scale observation studies using this technique may reveal unknown associations between treatments and mutational signatures, as well as new approaches to diagnostics.

In conclusion, we have demonstrated that restriction enzyme-based library preparation methods can be used to accurately recapitulate breast cancer mutation signatures and tumour mutation burden. Comparatively, RADseq-based methods offer improved TMB estimation when compared to a derived version of the widely used FoundationOne panel, which may aid in exploration of association between TMB and response to ICB. We also demonstrated the utility of restriction enzymes to characterise the genome of a model organism cell line and generation of episodic APOBEC mutagenesis in this cell line. Due to the relatively low cost and increased availability of these methods, they may be preferable to assays such as WGS or WES in a range of tumour types for estimation of TMB and mutation signature quantification where these parameters have prognostic/predictive value.

## Acknowledgements

We thank the members of the Centre for Translational Cancer Research, Dunedin for support during lab work. We also thank Shannon Clarke and Tracey Van Stijn from AgResearch New Zealand for assistance in generating RADseq libraries. Finally, we thank Vincent Caval from the Pasteur Institute for gifting the APOBEC encoding plasmids. This work was funded by grants from the New Zealand Breast Cancer Foundation.

## Supplementary figures

**Supplementary Figure 1.**
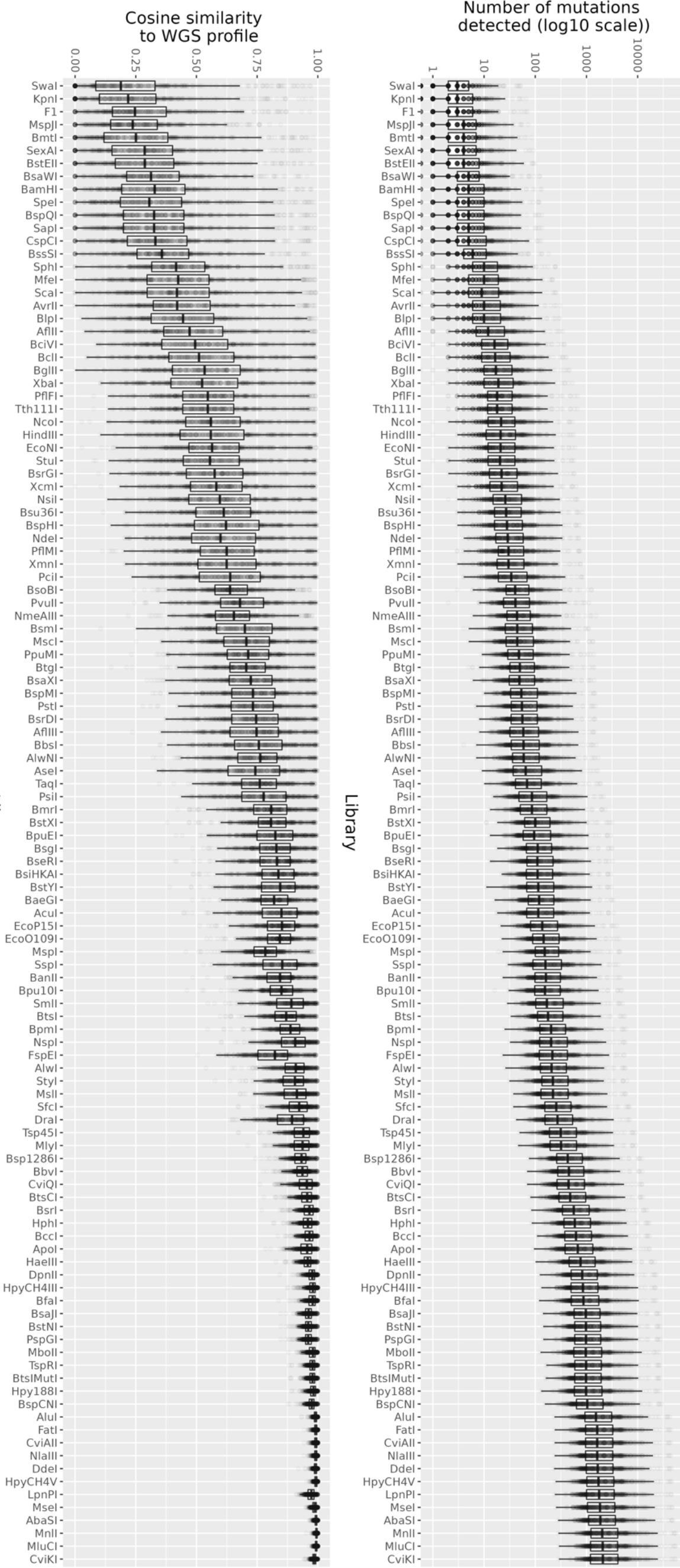
Extended library capture information. a) number of mutations captured by each library for each of 560 breast cancer genomes. b) Distribution of cosine similarity of the library 96 trinucleotide signature profile to the WGS trinucleotide signature profile.

**Supplementary Figure 2.**
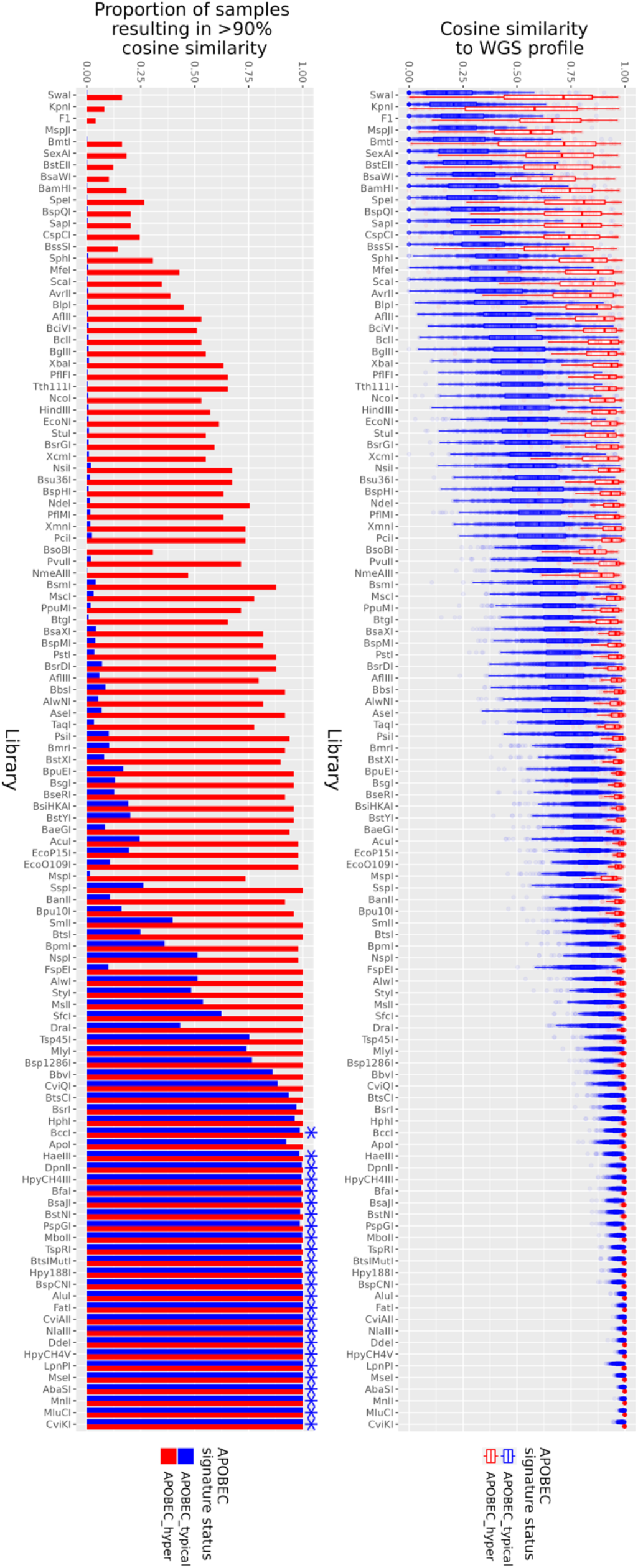
Mutation profile reconstruction by RADseq libraries. a) Distribution of cosine similarity of the library 96 trinucleotide signature profile to the WGS trinucleotide signature profile, split by APOBEC hypermutation status. d) Library vs proportion of profiles that result in 90% cosine similarity to WGS profile. Samples are coloured by APOBEC hypermutation status as in c).

**Supplementary Figure 3.**
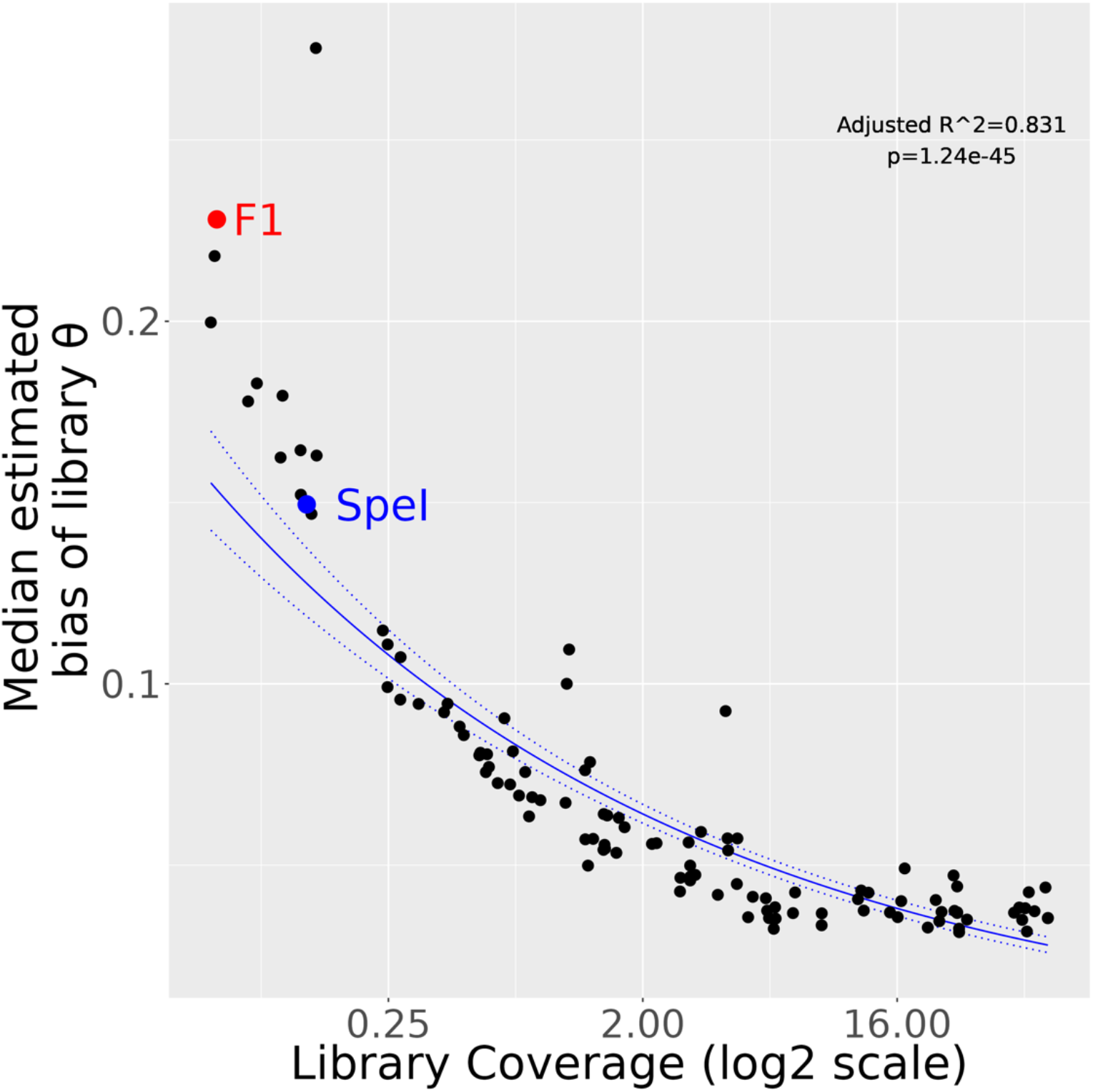
Estimated bias θ from Li, Luo (2021) of different libraries by library coverage. Each point represents a different library. F1 and SpeI libraries are highlighted as red and blue points respectively.

**Supplementary Figure 4.**
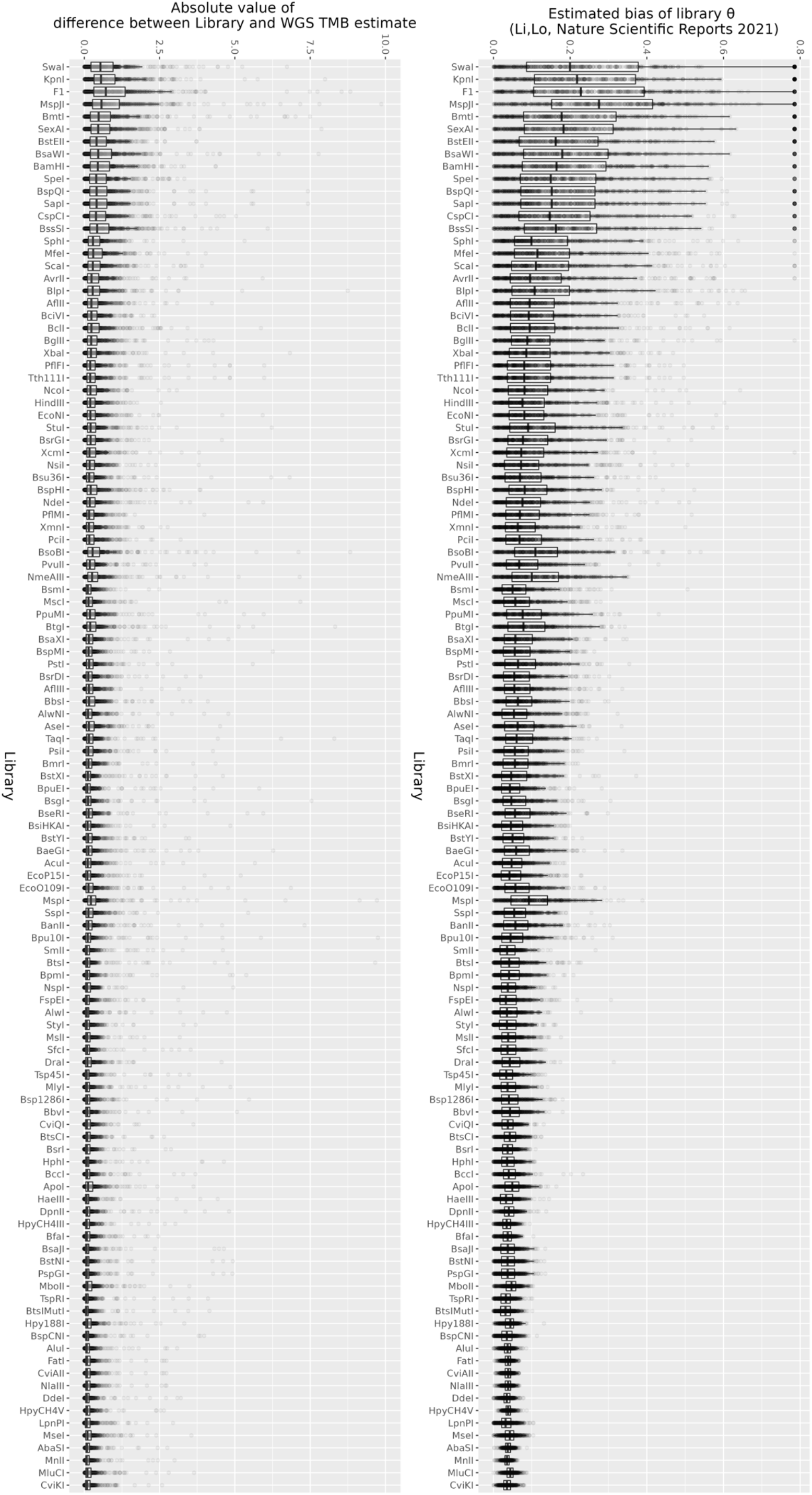
Measures of TMB estimation error in reduced representation libraries in 560 breast cancers. a) estimated bias of library θ (from Li, Luo, 2021) of each library. b) absolute value of difference between estimated TMB value from library and actual TMB value from WGS. In each plot, individual samples are represented as points.

**Supplementary Figure 5.**
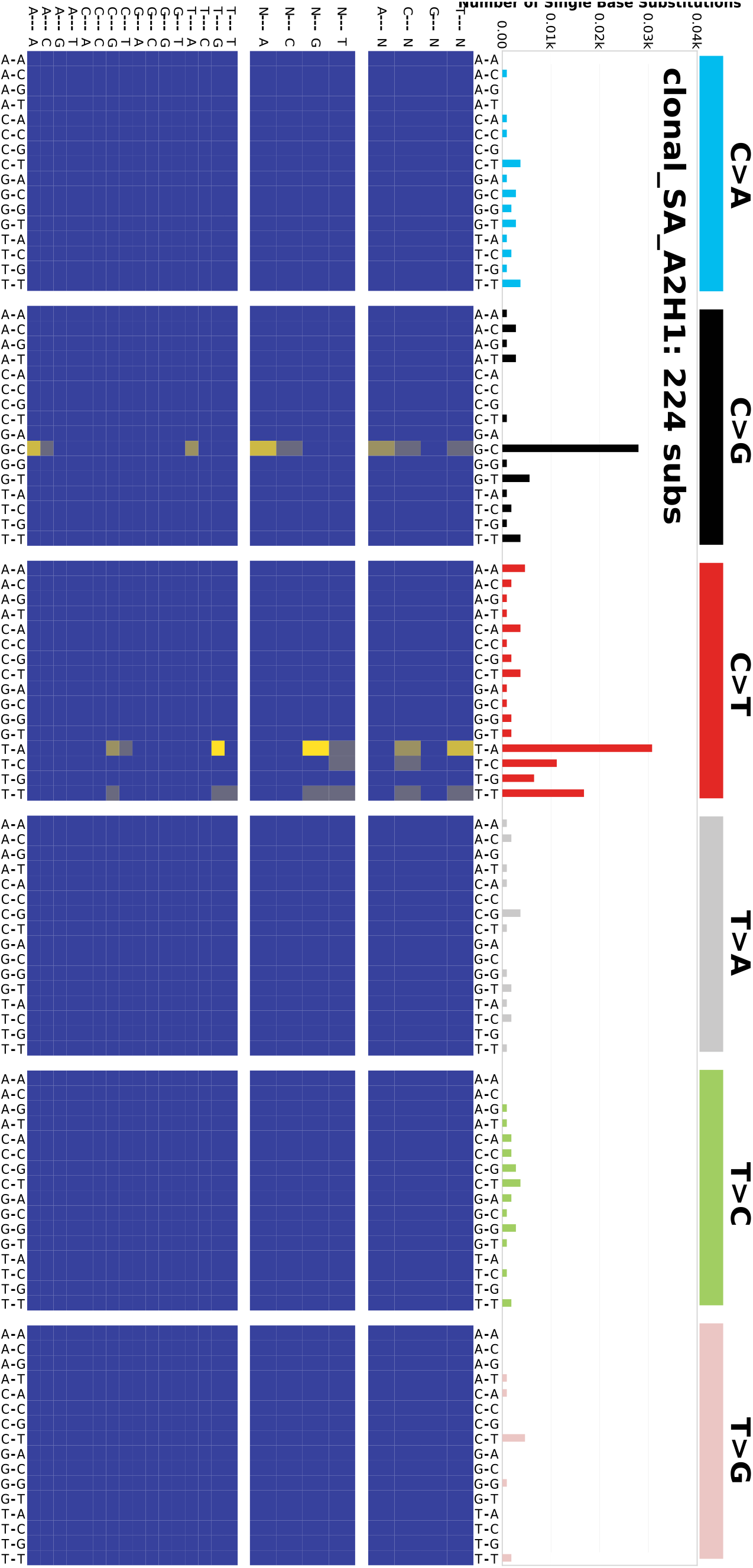
Extended 96 mutation profile of the A2H1 clone. Top: the proportion of mutations attributed to 96 mutation subtypes, coloured by mutation type and ordered by surrounding bases. Bottom: heatmap of relative contributions of mutations in extended contexts: the top heatmap is the extended context considering the bases 5’ of trinucleotide context; the middle heatmap is the extended context considering the bases 3’ of trinucleotide context; and the bottom heatmap is the extended context considering bases 5’ and 3’ of the trinucleotide context.

## References

1. Malone, E. R., Oliva, M., Sabatini, P. J. B., Stockley, T. L. & Siu, L. L. Molecular profiling for precision cancer therapies. Genome Med. 12, 8 (2020).

2. Subbiah, V., Solit, D. B., Chan, T. A. & Kurzrock, R. The FDA approval of pembrolizumab for adult and pediatric patients with tumor mutational burden (TMB) ≥10: a decision centered on empowering patients and their physicians. Ann. Oncol. 31, 1115–1118 (2020).

3. McGrail, D. J. et al. High tumor mutation burden fails to predict immune checkpoint blockade response across all cancer types. Ann. Oncol. 32, 661–672 (2021).

4. Alexandrov, L. B. et al. Signatures of mutational processes in human cancer. Nature 500, 415–421 (2013).

5. Alexandrov, L. B. et al. The repertoire of mutational signatures in human cancer. Nature 578, 94–101 (2020).

6. Burns, M. B. et al. APOBEC3B is an enzymatic source of mutation in breast cancer. Nature 494, 366–370 (2013).

7. Burns, M. B., Temiz, N. A. & Harris, R. S. Evidence for APOBEC3B mutagenesis in multiple human cancers. Nat. Genet. 45, 977–983 (2013).

8. Nik-Zainal, S. et al. Landscape of somatic mutations in 560 breast cancer whole-genome sequences. Nature 534, 47–54 (2016).

9. Barroso-Sousa, R. et al. Prevalence and mutational determinants of high tumor mutation burden in breast cancer. Ann. Oncol. 31, 387–394 (2020).

10. Tate, J. G. et al. COSMIC: the Catalogue Of Somatic Mutations In Cancer. Nucleic Acids Res. 47, D941–D947 (2019).

11. Davies, H. et al. HRDetect is a predictor of BRCA1 and BRCA2 deficiency based on mutational signatures. Nat. Med. 23, 517 (2017).

12. Wang, S., Jia, M., He, Z. & Liu, X.-S. APOBEC3B and APOBEC mutational signature as potential predictive markers for immunotherapy response in non-small cell lung cancer. Oncogene 37, 3924 (2018).

13. Boichard, A. et al. APOBEC-related mutagenesis and neo-peptide hydrophobicity: implications for response to immunotherapy. OncoImmunology 8, 1550341 (2019).

14. Boichard, A., Tsigelny, I. F. & Kurzrock, R. High expression of PD-1 ligands is associated with kataegis mutational signature and APOBEC3 alterations. Oncoimmunology 6, (2017).

15. Wu, H.-X., Wang, Z.-X., Zhao, Q., Wang, F. & Xu, R.-H. Designing gene panels for tumor mutational burden estimation: the need to shift from ‘correlation’ to ‘accuracy’. J. Immunother. Cancer 7, (2019).

16. Li, Y. & Luo, Y. Optimizing the evaluation of gene-targeted panels for tumor mutational burden estimation. Sci. Rep. 11, 21072 (2021).

17. Merino, D. M. et al. Establishing guidelines to harmonize tumor mutational burden (TMB): in silico assessment of variation in TMB quantification across diagnostic platforms: phase I of the Friends of Cancer Research TMB Harmonization Project. J. Immunother. Cancer 8, e000147 (2020).

18. Andrews, K. R., Good, J. M., Miller, M. R., Luikart, G. & Hohenlohe, P. A. Harnessing the power of RADseq for ecological and evolutionary genomics. Nat. Rev. Genet. 17, 81–92 (2016).

19. Perner, J. et al. The mutREAD method detects mutational signatures from low quantities of cancer DNA. Nat. Commun. 11, 3166 (2020).

20. R Core Team. R: A Language and Environment for Statistical Computing. (R Foundation for Statistical Computing, 2022).

21. Gehring, J. S., Fischer, B., Lawrence, M. & Huber, W. SomaticSignatures: inferring mutational signatures from single-nucleotide variants. Bioinformatics 31, 3673–3675 (2015).

22. Blokzijl, F., Janssen, R., van Boxtel, R. & Cuppen, E. MutationalPatterns: comprehensive genome-wide analysis of mutational processes. Genome Med. 10, 33 (2018).

23. Roberts, S. A. et al. An APOBEC Cytidine Deaminase Mutagenesis Pattern is Widespread in Human Cancers. Nat. Genet. 45, 970–976 (2013).

24. Milbury, C. A. et al. Clinical and analytical validation of FoundationOne®CDx, a comprehensive genomic profiling assay for solid tumors. PLoS ONE 17, e0264138 (2022).

25. Chan, S. R. et al. STAT1-deficient mice spontaneously develop estrogen receptor α-positive luminal mammary carcinomas. Breast Cancer Res. BCR 14, R16 (2012).

26. Caval, V., Suspène, R., Shapira, M., Vartanian, J.-P. & Wain-Hobson, S. A prevalent cancer susceptibility APOBEC3A hybrid allele bearing APOBEC3B 3′UTR enhances chromosomal DNA damage. Nat. Commun. 5, 5129 (2014).

27. Martin, A. S. et al. A panel of eGFP reporters for single base editing by APOBEC-Cas9 editosome complexes. Sci. Rep. 9, 497 (2019).

28. Schaefer, M. R. et al. A novel trafficking signal within the HLA-C cytoplasmic tail allows regulated expression upon differentiation of macrophages. J. Immunol. Baltim. Md 1950 180, 7804–7817 (2008).

29. Elshire, R. J. et al. A robust, simple genotyping-by-sequencing (GBS) approach for high diversity species. PLOS ONE 6, e19379 (2011).

30. Dodds, K. G. et al. Construction of relatedness matrices using genotyping-by-sequencing data. BMC Genomics 16, 1047 (2015).

31. Griffith, O. L. et al. Truncating Prolactin Receptor Mutations Promote Tumor Growth in Murine Estrogen Receptor-Alpha Mammary Carcinomas. Cell Rep. 17, 249–260 (2016).

32. Leinonen, R., Sugawara, H. & Shumway, M. The Sequence Read Archive. Nucleic Acids Res. 39, D19–D21 (2011).

33. Martin, M. Cutadapt removes adapter sequences from high-throughput sequencing reads. EMBnet.journal 17, 10–12 (2011).

34. Bolger, A. M., Lohse, M. & Usadel, B. Trimmomatic: a flexible trimmer for Illumina sequence data. Bioinformatics 30, 2114–2120 (2014).

35. Li, H. & Durbin, R. Fast and accurate short read alignment with Burrows-Wheeler transform. Bioinforma. Oxf. Engl. 25, 1754–1760 (2009).

36. Van der Auwera, G. A. & O’Connor, B. D. *Genomics in the Cloud*. (O’Reilly Media, Inc., 2020).

37. Li, Y. & Luo, Y. Optimizing the evaluation of gene-targeted panels for tumor mutational burden estimation. Sci. Rep. 11, 21072 (2021).

38. Giltnane, J. M. et al. Genomic profiling of ER+ breast cancers after short-term estrogen suppression reveals alterations associated with endocrine resistance. Sci. Transl. Med. 9, (2017).

39. Pan, J.-W. et al. Germline APOBEC3B deletion increases somatic hypermutation in Asian breast cancer that is associated with Her2 subtype, PIK3CA mutations, and immune activation. Int. J. Cancer (2021) doi:10.1002/ijc.33463.

40. Nik-Zainal, S. et al. Association of a germline copy number polymorphism of APOBEC3A and APOBEC3B with burden of putative APOBEC-dependent mutations in breast cancer. Nat. Genet. 46, 487–491 (2014).

41. Kidd, J. M., Newman, T. L., Tuzun, E., Kaul, R. & Eichler, E. E. Population stratification of a common APOBEC gene deletion polymorphism. PLoS Genet. 3, (2007).

42. Faden, D. L. et al. APOBEC mutagenesis is tightly linked to the immune landscape and immunotherapy biomarkers in head and neck squamous cell carcinoma. Oral Oncol. 96, 140–147 (2019).

43. Roper, N. et al. APOBEC mutagenesis and copy-number alterations are drivers of proteogenomic tumor evolution and heterogeneity in metastatic thoracic tumors. Cell Rep. 26, 2651–2666.e6 (2019).

44. Pan, J.-W. et al. Germline APOBEC3B deletion in Asian women increases somatic hypermutation in breast cancer that is associated with Her2 subtype, PIK3CA mutations, immune activation, and increased survival. *bioRxiv* 2020.06.04.135251 (2020) doi:10.1101/2020.06.04.135251.

45. Levinson, A. et al. Complete response to PD-1 inhibition in an adolescent with relapsed clear cell adenocarcinoma of the cervix predicted by neoepitope burden and APOBEC signature. *JCO Precis*. Oncol. 4, (2020).

46. Buchhalter, I. et al. Size matters: Dissecting key parameters for panel-based tumor mutational burden analysis. Int. J. Cancer 144, 848–858 (2019).

47. Zheng, M. Tumor mutation burden for predicting immune checkpoint blockade response: the more, the better. J. Immunother. Cancer 10, e003087 (2022).

48. Chalmers, Z. R. et al. Analysis of 100,000 human cancer genomes reveals the landscape of tumor mutational burden. Genome Med. 9, 34 (2017).

49. McGranahan, N. et al. Clonal neoantigens elicit T cell immunoreactivity and sensitivity to immune checkpoint blockade. Science 351, 1463–1469 (2016).

50. Nair, S., Sanchez-Martinez, S., Ji, X. & Rein, A. Biochemical and biological studies of mouse APOBEC3. J. Virol. 88, 3850–3860 (2014).

51. Petljak, M. et al. Mechanisms of APOBEC3 mutagenesis in human cancer cells. Nature 1–9 (2022) doi:10.1038/s41586-022-04972-y.

52. Perry, E. B. et al. Tumor diversity and evolution revealed through RADseq. Oncotarget 8, 41792–41805 (2017).

53. Zou, X. et al. Validating the concept of mutational signatures with isogenic cell models. Nat. Commun. 9, 1744 (2018).

