## Supplementary Figure 1 for "Restriction site associated DNA sequencing for tumour mutation burden estimation and mutation signature analysis"

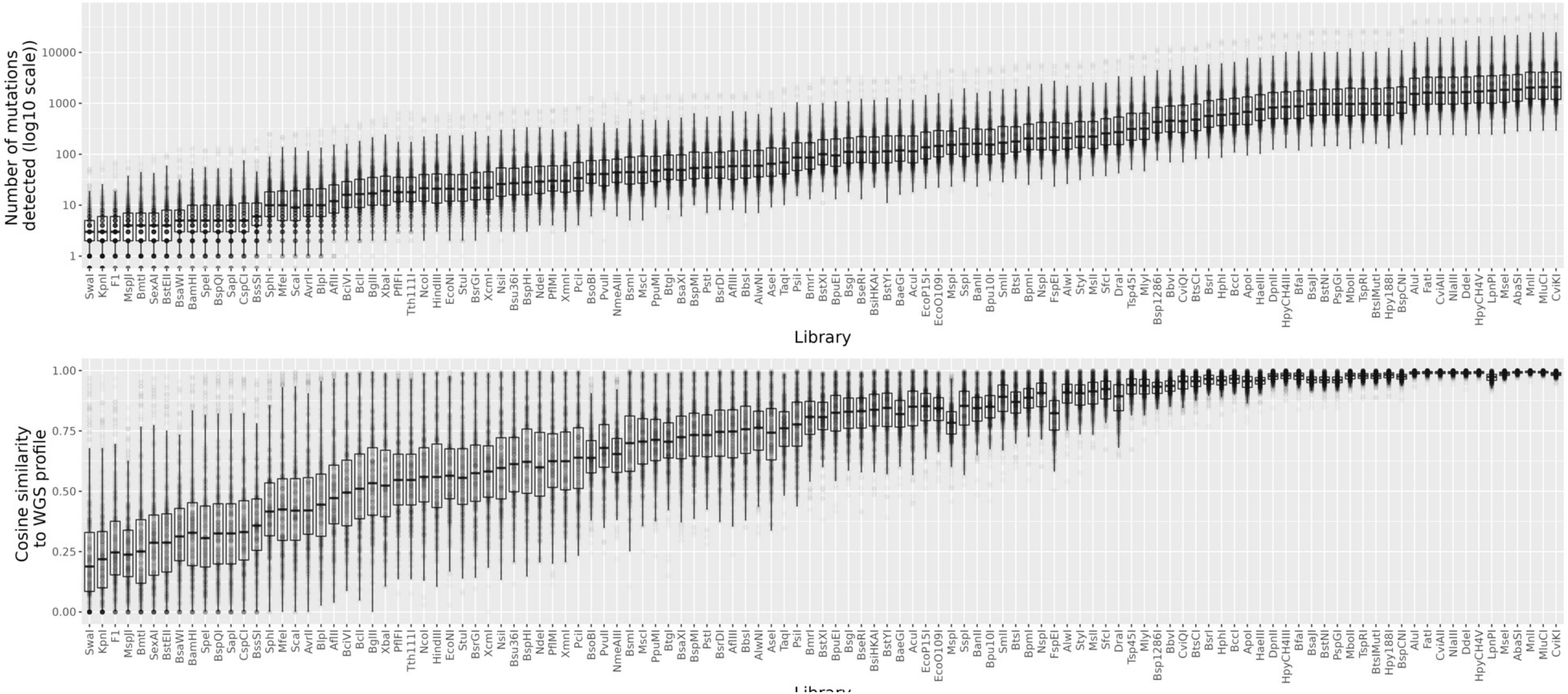


Supplementary Figure 1. Extended library capture information.

a) number of mutations captured by each library for each of 560 breast cancer genomes. b) Distribution of cosine similarity of the library 96 trinucleotide signature profile to the WGS trinucleotide signature profile.
