## Supplementary Figure 2 for "Restriction site associated DNA sequencing for tumour mutation burden estimation and mutation signature analysis"

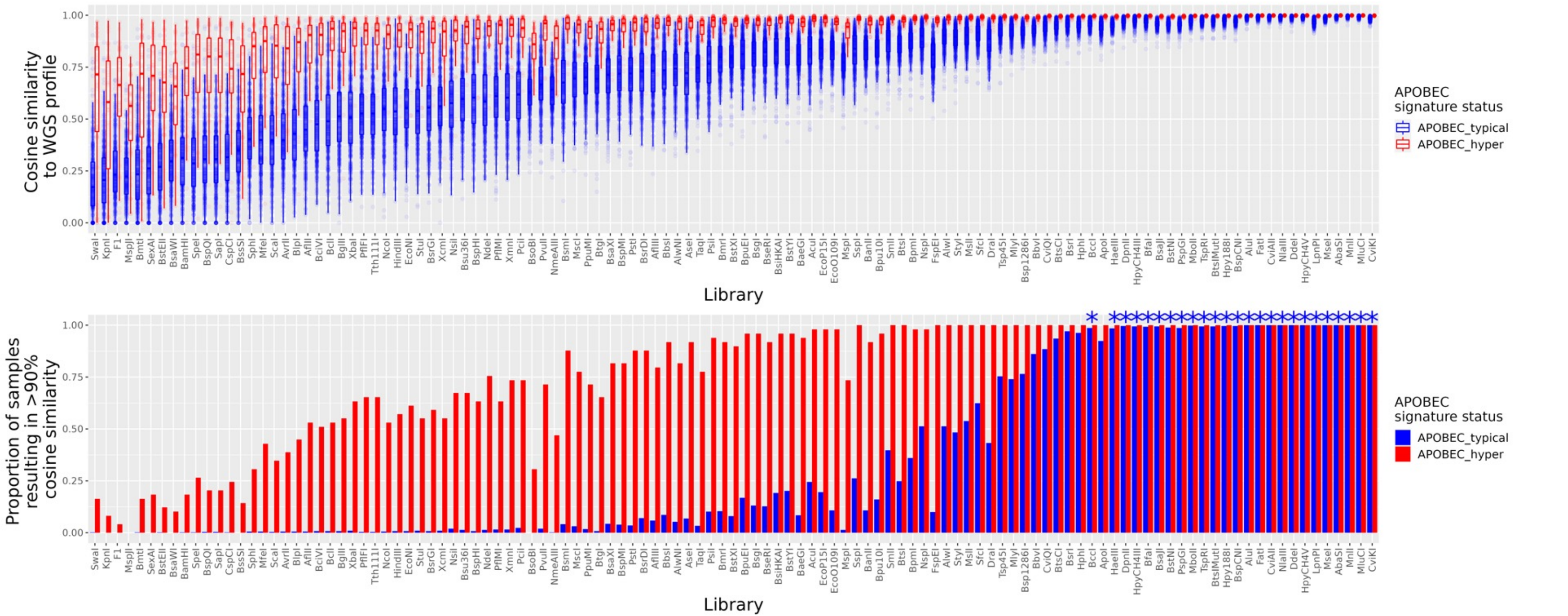


Supplementary Figure 2. Mutation profile reconstruction by RADseq libraries.

a) Distribution of cosine similarity of the library 96 trinucleotide signature profile to the WGS trinucleotide signature profile, split by APOBEC hypermutation status. d) Library vs proportion of profiles that result in 90% cosine similarity to WGS profile. Samples are coloured by APOBEC hypermutation status as in c).
