## Supplementary Figure 3 for "Restriction site associated DNA sequencing for tumour mutation burden estimation and mutation signature analysis"

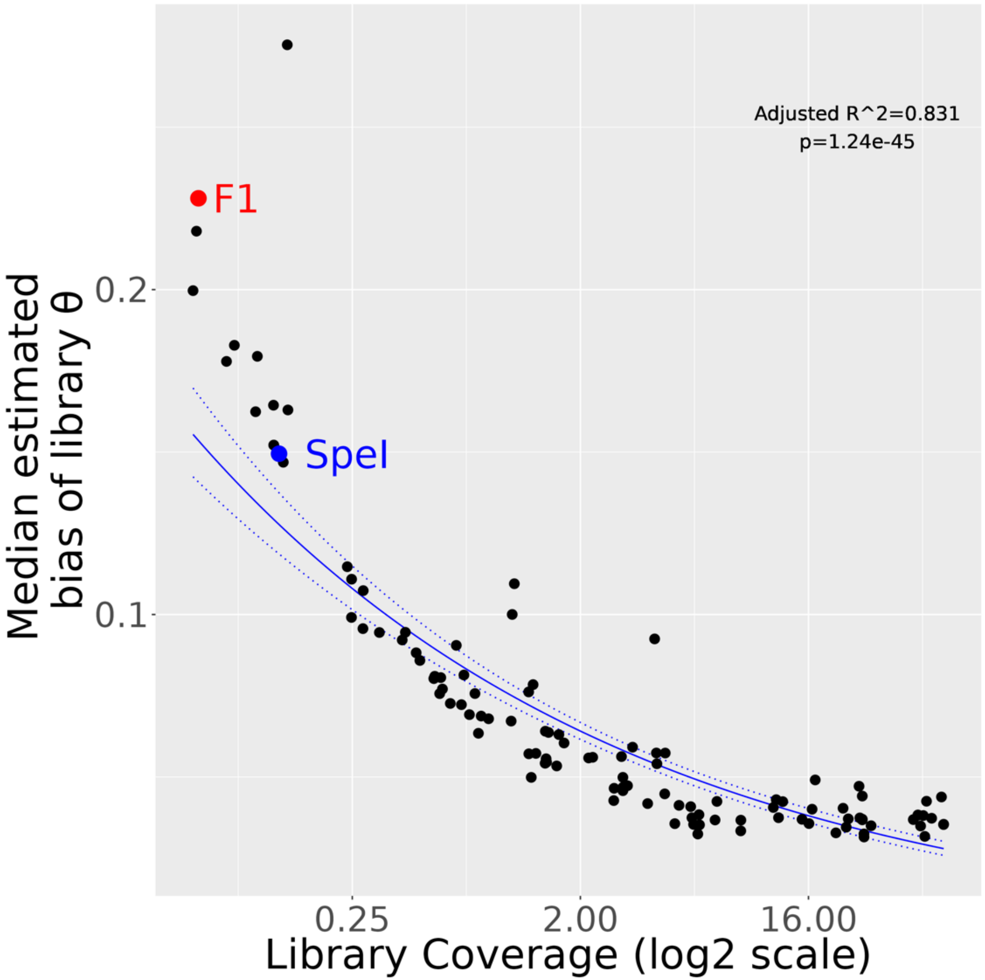


Supplementary Figure 3. Estimated bias θ from Li, Luo (2021) of different libraries by library coverage.

Each point represents a different library. F1 and SpeI libraries are highlighted as red and blue points respectively.
