## Supplementary Figure 4 for "Restriction site associated DNA sequencing for tumour mutation burden estimation and mutation signature analysis"

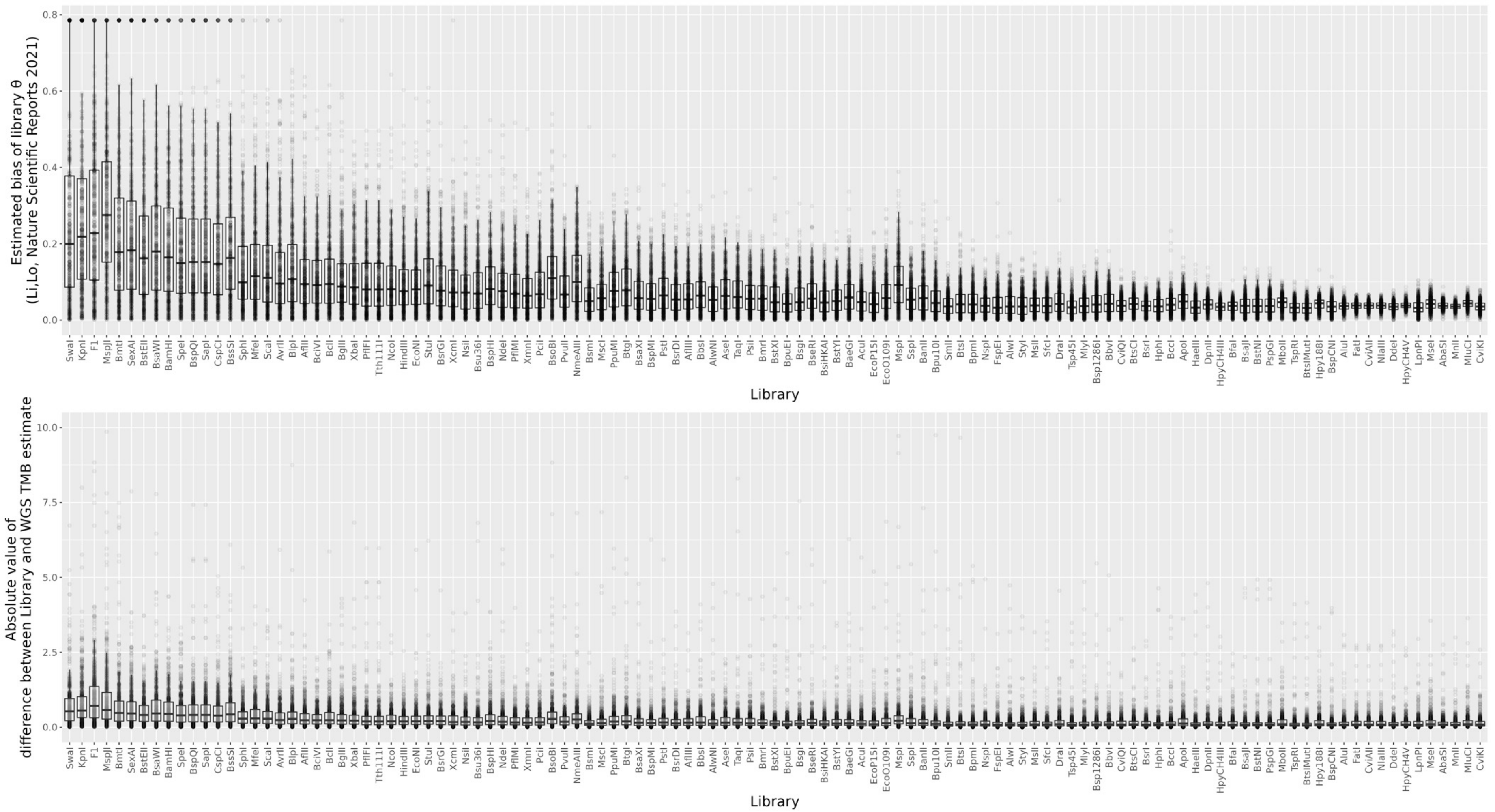


<u>Supplementary Figure 4  Measures of TMB estimation error in reduced representation libraries in 560 breast cancers.</u>

a) estimated bias of library θ (from Li, Luo, 2021) of each library. b) absolute value of difference between estimated TMB value from library and actual TMB value from WGS. In each plot, individual samples are represented as points.
