## Supplementary Figure 5 for "Restriction site associated DNA sequencing for tumour mutation burden estimation and mutation signature analysis"

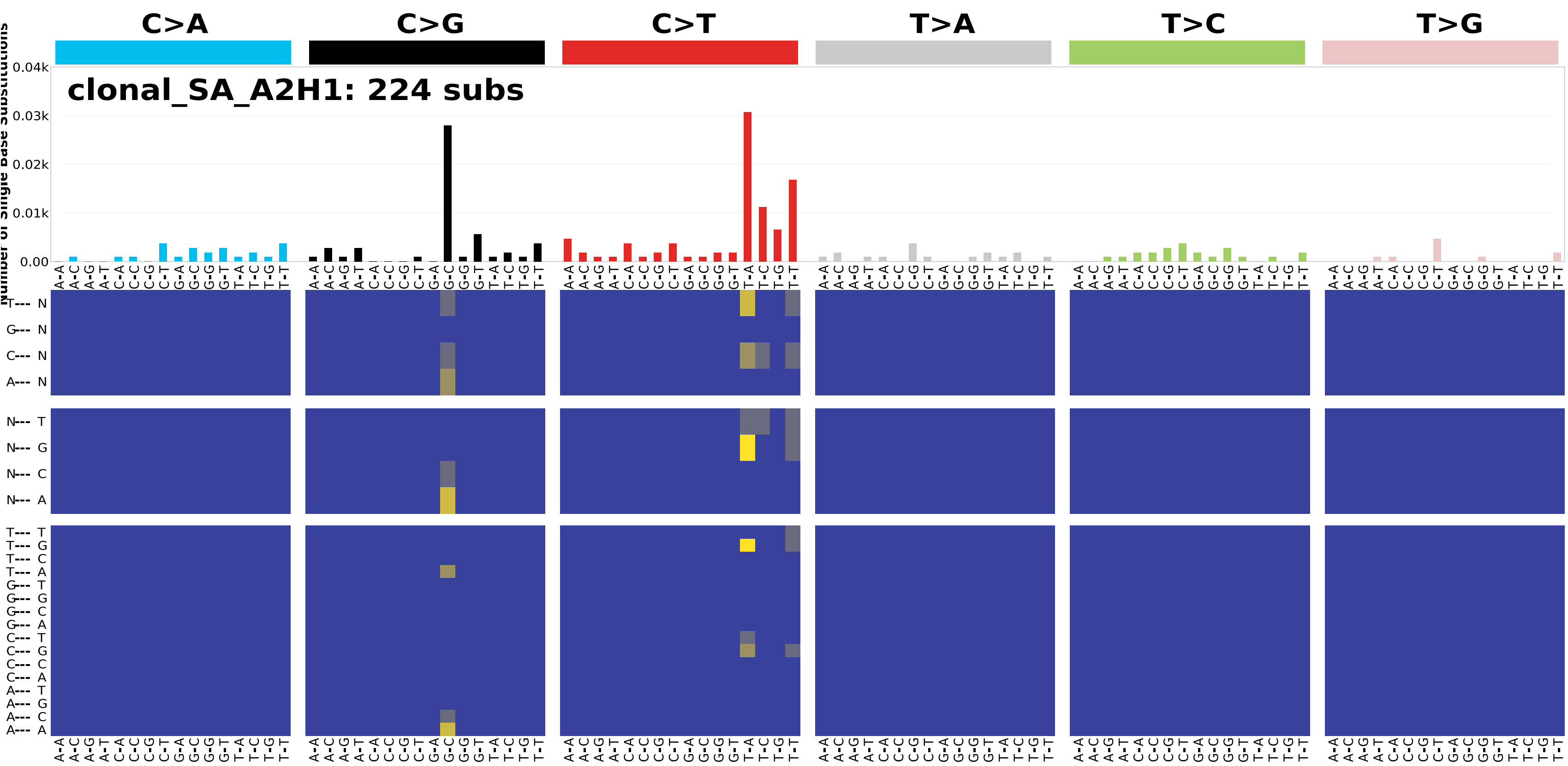


Supplementary Figure 5. Extended 96 mutation profile of the A2H1 clone.

Top: the proportion of mutations attributed to 96 mutation subtypes, coloured by mutation type and ordered by surrounding bases. Bottom: heatmap of relative contributions of mutations in extended contexts: the top heatmap is the extended context considering the bases 5’ of trinucleotide context; the middle heatmap is the extended context considering the bases 3’ of trinucleotide context; and the bottom heatmap is the extended context considering bases 5’ and 3’ of the trinucleotide context.
